# Tracing patterns of evolution and acquisition of drug resistant *Aspergillus fumigatus* infection from the environment using population genomics

**DOI:** 10.1101/2021.04.07.438821

**Authors:** Johanna Rhodes, Alireza Abdolrasouli, Katie Dunne, Thomas R. Sewell, Yuyi Zhang, Eloise Ballard, Amelie P. Brackin, Norman van Rhijn, Alexandra Tsitsopoulou, Raquel B. Posso, Sanjay H Chotirmall, Noel G McElvaney, Philip G Murphy, Alida Fe Talento, Julie Renwick, Paul S. Dyer, Adrien Szekely, Michael J. Bromley, Elizabeth M. Johnson, P. Lewis White, Adilia Warris, Richard C. Barton, Silke Schelenz, Thomas R. Rogers, Darius Armstrong-James, Matthew C. Fisher

**Author notes:** Corresponding authors: Johanna Rhodes –. Matthew Fisher –. Lee Kong Chian School of Medicine, Nanyang Technological University, Singapore (Sanjay H Chotirmall).

## Abstract

Infections caused by opportunistic fungal pathogens are increasingly resistant to first-line azole antifungal drugs. However, despite its clinical importance, little is known about the extent to which susceptible patients acquire infection from drug resistant genotypes in the environment. Here, we present a population genomic analysis of the mould *Aspergillus fumigatus* from across the United Kingdom and Republic of Ireland. First, we show occurrences where azole resistant isolates of near identical genotypes were obtained from both environmental and clinical sources, indicating with high confidence the infection of patients with resistant isolates transmitted from the environment. Second, we find that the fungus is structured into two clades (‘A’ and ‘B’) with little interclade recombination and the majority of environmental azole resistance genetically clustered inside Clade A. Genome-scans show the impact of selective sweeps across multiple regions of the genome. These signatures of positive selection are seen in regions containing canonical genes encoding fungicide resistance in the ergosterol biosynthetic pathway, whilst other regions under selection have no defined function. Phenotyping identified genes in these regions that could act as modifiers of resistance showing the utility of reverse genetic approaches to dissect the complex genomic architecture of fungal drug resistance. Understanding the environmental drivers and genetic basis of evolving fungal drug resistance needs urgent attention, especially in light of increasing numbers of patients with severe viral respiratory tract infections who are susceptible to opportunistic fungal superinfections.

## Introduction

Fungal infections are often neglected, yet affect more than a billion people worldwide; mortality caused by fungal diseases matches that found for malaria or tuberculosis^1^. *Aspergillus fumigatus* is a globally ubiquitous environmental mould, which may opportunistically cause fungal lung disease. Although healthy human lungs clear inhaled inocula, invasive aspergillosis (IA) can occur in the ever increasing at-risk population with severe neutropenia, haematopoietic stem cell or solid organ transplants, those receiving immunosuppressive therapy (e.g. autoimmune diseases), and is emerging as an important pathogen as an influenza- and COVID-19 associated infection^3, 22^. Patients who suffer chronic infection by *A. fumigatus* include those with pulmonary conditions such as cystic fibrosis (CF) and severe asthma. CF patients are particularly at high risk, with 30% developing *Aspergillus*-related bronchitis and 18% developing allergic bronchopulmonary aspergillosis (ABPA) in adulthood^4^. Collectively, millions of individuals suffer from aspergillosis worldwide, with over 2.25 million occurring in the European Union alone^5^ testifying to the broad scale nature of the problem.

Triazole antifungal drugs are usually effective against *A. fumigatus* and comprise firstline therapy for prophylaxis and treatment of IA^6^. However, emerging resistance to azoles has been reported^7^ worldwide for both environmental and clinical isolates^8, 9^. Triazole antifungal drug resistance has serious clinical implications, with retrospective studies of patients infected with IA-associated azole-resistant *A. fumigatus* showing a 25% increase in mortality at day 90 when compared with patients with wild-type-infections^10^. While *in vivo* emergence of resistance during extended azole therapy is well documented^11, 12^, *ex vivo* evolution of resistance in the environment as a result of exposure to sterol 14α-demethylation inhibitor (DMI) fungicides has also been postulated^13, 14^. These agricultural chemicals have been increasingly and widely used since their development in the 1970’s and affect pan-fungal sterol and related membrane biosynthesis by targeting the enzyme 14α-sterol demethylase. Azole resistance in *A. fumigatus* is thought to occur *via* the unintended selection by DMIs for isolates with reduced sensitivity due to polymorphisms in the *cyp51A* gene that encodes this membrane-building enzyme. Broadly, azole-resistance in *A. fumigatus* occurring in the environment is characterised by signature mechanisms involving expression-upregulating tandem repeats (TR) in the promoter region of *cyp51A* accompanied by point mutations within this gene which decrease the affinity of azoles for the target protein; the most commonly occurring alleles are known as TR_34_/L98H and TR_46_/Y121F/T289A and are associated with high level itraconazole and voriconazole resistance respectively both inside and outside of the clinic^15–17^. The spatially widespread occurrence of these canonical alleles alongside increasing reports of more complex *cyp51A* resistance-associated polymorphisms^18–20^ underpin the hypothesis that the broad application of agricultural azole fungicides is driving natural selection, amplification and ultimately acquisition of azole-resistant airborne *A. fumigatus* conidia by susceptible patients^21^. Furthermore, the potential for global spread of these resistance mechanisms through floriculture products, especially plant bulbs, has been demonstrated^22^, while the global dispersal of conidia on air-currents is impossible to contain.

There is a need to explore the potential link between the increasing clinical incidence of azole-resistant aspergillosis and the increasingly broad range of azole resistant genotypes that are being reported in the environment^23^ using modern genomic epidemiological methods. The rate at which environmental resistance develops will be determined by the strength of natural selection by azole fungicides acting on beneficial mutations, a process that is further influenced by recombination, gene flow, and dispersal with the latter likely being substantial for *A. fumigatus* given its ubiquitous presence in nature and airborne buoyancy of conidia. Whilst azole-resistant alleles can potentially segregate into diverse genetic backgrounds through sex^24^, the majority of *A. fumigatus* reproduction in the environment is thought to occur asexually^26^ producing clones with high genetic similarity. Therefore, *a priori* expectations are that the genomes of azole-resistant *A. fumigatus* will exhibit the genetic signatures of linkage, selective sweeps, and divergence of selected loci from the wild-type population that is predicated on the amount of population-wide recombination. Ultimately, clonality will cause high genetic similarity between clinical isolates and their near-contemporaneous environmental progenitors, discovery of which constitutes epidemiological evidence for the acquisition of azole resistant aspergillosis. Evidence for these expectations comes from a recent global study by our laboratory demonstrating the non-random distribution of azole resistance in *A. fumigatus* multilocus microsatellite genotypes worldwide, compatible with selective sweeps caused by genetic hitchhiking alongside azole-resistance polymorphisms^27^.

Here, we used whole genome sequencing to interrogate the spatial and molecular epidemiology and population genomics of a broad panel of *A. fumigatus* isolates collected from medical centres and environmental sites across England, Wales, Scotland and the Republic of Ireland. The aim of our study was to understand the genetic architecture of *A. fumigatus* populations in this region in order to prove, or disprove, the proposed link between azole-resistance found in the environment and patients.

## Methods

### Fungal isolates

A total of 218 isolates were included in this study. 153 clinical *A. fumigatus* isolates from five participating centres were included. Patients either had respiratory colonisation with *A. fumigatus* or were suffering from different manifestations of aspergillosis and had the following underlying disorders (Table S1): cystic fibrosis (64%), other conditions (11%), unknown (25%). Environmental isolates (*n* = 65) were collected as described in Sewell *et al.*, Dunne *et al.*, and Tsitsopoulou *et al.*^22, 28, 29^, and collected from the following sources: soil (72%), plant bulbs (12%), air (3%), compost (2%), and unknown (11%). Many isolates from both clinical and environmental sources were specifically selected for whole genome sequencing because they displayed phenotypic azole resistance (raised minimum inhibitory concentrations (MICs) to at least one triazole drug using EUCAST or CLSI) and do not constitute a randomised sample.

**Table 1:** Relative frequencies of metadata associated with Clades A and B within this dataset

|  | Clade A (total 123 isolates) | Clade B (total 95 isolates) |
| --- | --- | --- |
| Number of clinical isolates | 71 (58%) | 82 (86%) |
| Number of environmental isolates | 52 (42%) | 13 (14%) |
| *MAT1-1* | 48 (39%) | 31 (33%) |
| *MAT1-2* | 75 (61%) | 64 (66%) |
| *Cyp51A* wildtype | 24 (20%) | 82 (86%) |
| *Cyp51A* AMR polymorphisms | 99 (80%) | 13 (14%) |

All referred isolates were identified by phenotypic morphology as *A. fumigatus* species complex based on colonial morphology and microscopic characteristics. Isolates were cultured on Sabouraud dextrose agar (Oxoid, Basingstoke, UK) and malt extract agar (Sigma Aldrich) at 37°C ± 2°C for 5-7 days in the dark. Adhesive tape technique was used for microscopic examination of fungal cultures. Slides were prepared with lactophenol-cotton blue as mounting and staining fluid. The Atlas of Clinical Fungi (https://www.clinicalfungi.org) was consulted as an identification reference. In addition, growth at 45°C was used to exclude most cryptic species within section *Fumigati*. Isolates with elevated azole MICs were confirmed to be *A. fumigatus* by matrix-assisted laser desorption ionization–time of flight mass spectrometry (MALDI–TOF MS), performed with a Microflex LT system (Bruker Daltonics, Bremen, Germany) using Biotyper 3.0 software with the additional fungi library (Bruker Daltonics, Bremen, Germany) according to the manufacturer’s recommendations, or as described by Dunne *et al.*^22^. Exact identification of azole-resistant *A. fumigatus* isolates from two participating centres in London, UK, was confirmed by sequencing the partial calmodulin gene (*CaM* locus) as previously described^30^. Antifungal susceptibility testing was completed as part of the original sampling studies, or determined according to the standard EUCAST^31^ or CLSI M38-A2 broth microdilution methods^32^. MICs for itraconazole and voriconazole were determined for 92% of isolates (*n* = 200 and 201 respectively). MICs for posaconazole were determined for 81% of isolates (*n* = 177). Seventeen isolates (8%) were not tested for susceptibility and therefore have no recorded MICs for any antifungal drug.

### DNA preparation and whole-genome sequencing

High molecular weight DNA was extracted from all 218 isolates and quantified as described in Abdolrasouli *et al.*^21^. Genomic DNA libraries were constructed with the Illumina TruSeq Nano kit (Illumina, San Diego, CA) at NERC Biomolecular Analysis Facility (NBAF), University of Edinburgh, Scotland, UK (http://genomics.ed.ac.uk/). Prepared whole-genome libraries were sequenced on an Illumina HiSeq 2500 sequencer at NBAF, generating 150-bp paired end reads in high output mode. All raw reads have been submitted to the European Nucleotide Archive (ENA) under project accession number PRJEB27135. Six isolates (ARAF001-6) were sequenced as part of Abdolrasouli *et al.*^21^, with reads deposited under project accession number PRJEB8623.

### Bioinformatic analyses

All raw Illumina paired-end reads were quality checked using FastQC (v0.11.5; Babraham Institute) and aligned to the reference genome Af293^33^ using Burrows-Wheeler Aligner (BWA v0.7.8)^34^ mem and converted to sorted BAM format using SAMtools v1.3.1^35^. Variant calling was performed using GATK^36, 37^ HaplotypeCaller v3.6, excluding repetitive regions identified using RepeatMaster^38^ v4.0.6. Low confidence variants were filtered out providing they met at least one of the parameters “DP < 10 || MQ < 40.0 || QD < 2.0 || FS > 60.0 || ABHom < 0.9”. All variant calls with a minimum genotype quality of less than 50 were also removed using a custom script. Single nucleotide polymorphisms (SNPs) were mapped to genes using VCF-annotator (Broad Institute, Cambridge, MA).

### Phylogenetic analysis

Whole genome single nucleotide polymorphism (SNP) data were converted into presence/absence of a SNP with respect to the reference. SNPs identified as low confidence in the variant filtration step were converted to missing data. These binary data were converted into relaxed interleaved Phylip format, and maximum-likelihood phylogenies were constructed to assess sequence similarity between isolates using rapid bootstrap analysis over 1000 replicates using the BINCAT model of rate heterogeneity in RAxML^39^ v8.2.9. Phylogenies were visualised in FigTree v1.4.2.

### Analysis of genetic diversity and population inference

Genetic similarity and population allocation was investigated *via* hypothesis free approaches. Principal Component Analysis (PCA) and Discriminant Analysis of Principal Components (DAPC)^40^ was performed to interrogate the relationship between clinical and environmental isolates based on SNP data using the R package adegenet^41^ version 2.1.1. Genetic clusters were allocated based on identifying the optimal number of clusters (*k*) corresponding to the lowest Bayesian Information Criterion (BIC). STRUCTURE v2.3.4^42^ was used to independently investigate the population structure assignments that were predicted by DAPC and PCA.

We analysed a coancestry matrix based on whole-genome SNP data to assign individuals to populations *via* model-based clustering using *fineStructure*^43^ v2.0.7. *FineStructure* uses chromosome painting to identify important haplotypes and describe shared ancestry within a recombining population. These analyses were performed using a pan-clade genome-wide SNP matrix to infer recombination, population structure and assignment, and ancestral relationships of all lineages. ChromoPainter reduced the SNP matrix to a pairwise similarity matrix under a linked model, using information on linkage decay and reducing the within-population variance of the coancestry matrix relative to the between-population variance. Sliding-window population genetic statistics (Tajima’s *D*, nucleotide diversity π and *F*_ST_) were calculated for non-overlapping windows of 10 kb using vcftools^44^ v0.1.13, with resulting graphics produced in R v3.5.3 using ggplot2.

The index of association and rBarD are commonly used to estimate linkage disequilibrium. These two statistics were calculated using Poppr v2.8.5^45^ in R v3.5.3 using 999 resamplings of the data under the null hypothesis of recombination.

### Identifying loci associated with itraconazole resistance using genome-wide association

Loci associated with itraconazole resistance were identified using *treeWAS*^46^, a method that was recently developed to address challenges specific to microbial genome-wide association studies (GWAS). *TreeWAS* uses phylogenetic information to correct for the microbial population structure; we therefore used the phylogeny presented in this study along with a nucleotide alignment for all 218 isolates. *TreeWAS* was performed for all isolates with MIC information with a *p*-value cut-off of 0.01 for three tests of association (subsequent, simultaneous and terminal) between azole susceptibility phenotype and genotype. The phenotype information was encoded as a discrete vector based on above the MIC breakpoint for itraconazole (and therefore resistant) or below (susceptible). Significant SNPs common to all three tests were combined and were mapped to their respective genes *via* FungiDB (release 50 *beta*)^47^.

### Drug sensitivity screening

Null mutants were obtained from the COFUN genome wide knockout collection^48^ and the COFUN transcription factor knockout library^49^. MFIG001 (A1160p+) was used as the parental isolate^50^. Strains were inoculated in 25 cm^2^ tissue culture flasks containing Sabouraud Dextrose Agar (Oxoid, Basingstoke, UK) + 100 µM hygromycin and cultured for 3 days at 37 °C. Conidia were harvested in phosphate buffered saline (PBS) + 0.01% Tween-20 (Sigma Aldrich, UK) by filtration through Miracloth (VWR, UK). Spore concentrations were determined by haemocytometer. Spores were inoculated in a CytoOne 96-well plate (StarLab, Milton Keynes, UK) containing RPMI-1640 medium 2% glucose and 165 mM MOPS buffer (pH 7.0) with 16-0.06 mg/L itraconazole. MICs were determined visually after 48 hours as outlined by EUCAST^51^. Each strain was assessed in biological triplicate. Heatmaps were generated in Graphpad PRISM (version 9).

## Results

### Whole genome sequencing of 218 isolates

Our collection spans a period of 12 years (2005-2017), covering a spatial range of 63,497 miles^2^ in England, Wales, Scotland and Republic of Ireland (Figure 1). Whole genome sequencing of 218 isolates of *A. fumigatus* isolated in England, Wales, Scotland and Republic of Ireland was performed (Table S1).

**Figure 1.**
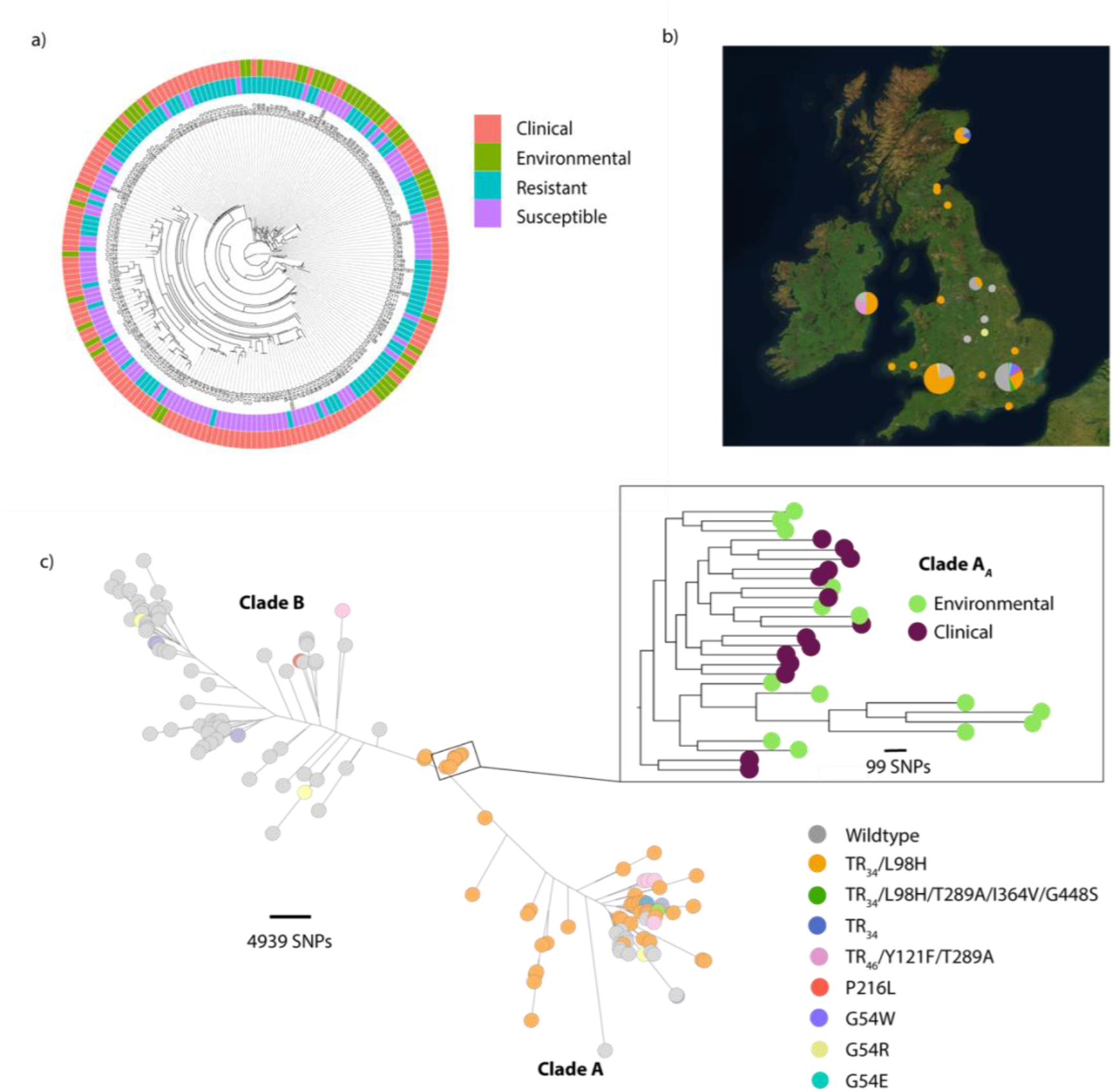
Phylogeographic and phenotypic variation of 218 *A. fumigatus* isolates in the United Kingdom and Republic of Ireland. a) Unrooted maximum likelihood phylogenetic tree (constructed in RAxML using genome-wide SNPs) showing the itraconazole MIC breakpoint (defined as above or below 2 mg/L for resistance or susceptibility, respectively) and clinical or environmental source of isolation. b) Map showing the location of isolation for isolates included in this study, with the legend (bottom right) indicating *cyp51A* polymorphisms present. c) Unrooted maximum likelihood phylogeny of all 218 isolates showing the genetic relationship between the isolates. ‘Clade A’ and ‘Clade B’ indicate the clustered nature of triazole resistance polymorphisms. The subclade in the midpoint of the phylogeny indicates a clonal clade, Clade A*_A_*, that is rich in clinical and environmental *A. fumigatus* isolates which contain the drug resistance polymorphism TR_34_/L98H, highlighted in the inset phylogeny.

Of these 218 sequenced isolates, 153 (70%) were clinical in origin, and the remaining 65 (30%) originated from environmental sources in the UK and Ireland. All reads mapped to >93.4% of the Af293 reference genome with an average 38x coverage (Supplementary Information Table 1). Within this dataset, just over a third of isolates (34%; *n* = 79) were found to contain the *MAT1-1* mating type idiomorph (Table S2). Mating type was reported for all isolates; a chi-square test showed a significant bias towards the *MAT1-2* idiomorph (*P* < 4.82983e^-05^), primarily seen in the environmental (*P* < 1.00183e^-05^), and to a lesser extent the clinical (*P* < 0.04396), populations (Table S2). The genomic dataset can be browsed as a Microreact project^52^ at https://microreact.org/project/6KR8996ywtVRV5wm233YhP (Figure S1).

**Table 2:** Relative frequencies of *cyp51A* genotypes and mating-type idiomorphs for clinical and environmental *A. fumigatus* isolates in this study

| *Cyp51A* genotype | Number of clinical isolates<br>(% of clinical isolates) | Number of environmental<br>isolates (% of<br>environmental isolates) |
| --- | --- | --- |
| TR<sub>34</sub>/L98H | 44 (28.8%) | 41 (63.1%) |
| TR<sub>34</sub> | 0 | 1 (1.5%) |
| TR<sub>34</sub>/L98H/T289A/I364V/G448S | 4 (2.6%) | 0 |
| TR<sub>46</sub>/Y121F/T289A | 0 | 7 (10.8%) |
| P216L | 2 (1.3%) | 0 |
| G54W | 9 (5.9%) | 0 |
| G54E | 1 (0.6%) | 0 |
| G54R | 2 (1.3%) | 1 (1.5%) |
| Wildtype | 91 (59.5%) | 15 (23.1%) |
| *MAT1-1* | 64 (41.8%) | 15 (23.1%) |
| *MAT1-2* | 89 (56.2%) | 50 (76.9%) |

EUCAST clinical antifungal breakpoints (www.eucast.org) were used to determine the susceptibility of isolates, with the exception of Irish isolates, where susceptibility was determined using CLSI methodology as described by Dunne *et al*.^22^. As clinical breakpoints from EUCAST and CLSI were broadly similar for *A. fumigatus* they were pooled in our analyses. Other studies have also found CLSI and EUCAST MICs to be similar^53, 54^. We report (Table S3) minimum inhibitory concentrations (MICs) for itraconazole (*n* = 200), voriconazole (*n* = 201) and posaconazole (*n* = 178). Of the isolates tested, 106 (49%) showed resistance to at least one of the test antifungal drugs. Regarding specific azoles, 48% (*n* = 96) exceeded MIC breakpoints to itraconazole (≥2 mg/L), 29% (*n* = 64) to voriconazole (≥2 mg/L), and 21% (*n* = 44) to posaconazole (≥0.5 mg/L). We found 26 isolates (12%) that exceeded MIC breakpoints to two or more azole drugs from both clinical (*n* = 23) and environmental (*n* = 3) sources. Seventeen isolates were not tested for antifungal susceptibility *via* EUCAST or CLSI; of these, 14 contained drug resistance polymorphisms (TR_34_/L98H or TR_46_/Y121F/T289A) and three contained no known drug resistance polymorphism. Thirteen isolates reported raised MICs but displayed no known drug resistance polymorphisms. These isolates are summarised in Table 3.

**Table 3:** *A. fumigatus* isolates presented in this study with raised MICs to antifungal azole drugs, yet with no known resistance polymorphism within *cyp51A*. ND = not done; Minimum Inhibitory Concentrations above EUCAST (2018) clinical breakpoints, and therefore resistant, to itraconazole (ITC), voriconazole (VOR) and Posaconazole (POS) are shaded in grey.

| Isolate | ITC MIC (mg/litre) | VOR MIC (mg/litre) | POS MIC (mg/litre) |
| --- | --- | --- | --- |
| C117 | 1 | 2 | ND |
| C118 | 8 | 1 | ND |
| C127 | >16 | 0.25 | 0.125 |
| C128 | >16 | 2 | 1 |
| C129 | 8 | 2 | 0.5 |
| C136 | >16 | 2 | 2 |
| C137 | >16 | 2 | 2 |
| C139 | >16 | 2 | 1 |
| C148 | 2 | 0.125 | 0.03 |
| C162 | 2 | 2 | 1 |
| C164 | 2 | 1 | 0.125 |
| C165 | 2 | 0.5 | 0.125 |
| C168 | 2 | 0.25 | 0.03 |

Among the 218 genomes, 329,261 sites along the 30 Mb genome were found to be polymorphic (∼1.1%), similar to that seen in our previous study using a small number of isolates recovered across a global-scale^21^. Pairwise identities show that, on average, each isolate of *A. fumigatus* in this dataset differs from all others by 11,828 single nucleotide polymorphisms (SNPs) testifying to the highly-discriminatory nature of this genotyping method. Tandem repeats and SNPs causing non-synonymous amino acid substitutions in *cyp51A* encoding 14-α lanesterol demethylase, the target of triazole antifungals, are summarised in Table S1 for all isolates where present. Identified polymorphisms were compared to known polymorphisms conferring resistance using the MARDy database^55^. The majority of clinical isolates (n = 91 (59.5%)) exhibited wildtype *cyp51A*, whilst TR_34_/L98H was predominant in environmental (*n* = 41 (63.1%)) isolates. The TR_34_/L98H polymorphism was the drug resistance allele found to occur with highest frequency in both clinical (*n* = 44 (28.8%)) and environmental populations (*n* = 41 (63.1%)) respectively; a second published polymorphism, TR_46_/Y121F/T289A, was only found in the environmental population (*n* = 7). Other *cyp51A* nonsynonymous polymorphisms known to be associated with azole resistance were also present within this dataset including: P216L (*n* = 2), G54W (*n* = 9), G54E (*n* = 1), G54R (*n* = 3) (Figure 1). These genotypes and their relative frequencies are presented in Table 2. The TR_46_/Y121F/T289A genotype and a single TR_34_ only (without the L98H substitution) were only recovered from environmental isolates. Alongside known resistance polymorphisms we found a novel *cyp51A* genotype associated with azole-resistance within the UK, the TR-associated polymorphism, TR_34_/L98H/T289A/I364V/G448S, which was detected in four isolates (C155-C158) collected in 2016 from a sputum sample of one patient with necrotising aspergillosis; these four isolates were found to contain this novel polymorphism, manifesting a multi-azole resistant phenotype (Table S2), demonstrated by high MIC values for itraconazole, voriconazole (both ≥ 16 mg/L) and posaconazole (4 mg/L). These isolates (with the exception of C155) were phenotypically distinct from classical *A. fumigatus* displaying slower growth rates with less pigmentation and fewer or no conidia compared to classical *A. fumigatus*, yet were confirmed to be *A. fumigatus* by mass spectrometry and the subsequent genome sequencing. All four isolates were *MAT1-2*, and were only separated by 145 SNPs on average, compared to over 11,000 SNPs that on average separate any pair of isolates in this dataset. This allele was only recovered in-patient and was not found in the environment (Table 2). Known non-TR-associated resistance markers P216L, G54W and G54E were only recovered in clinical isolates in this study despite having been isolated from environmental samples in previous studies^25, 56, 57^ (Table 2).

A single environmental isolate (C365) was found to contain only the 34-bp tandem repeat in the *cyp51A* promoter region, uncoupled from L98H. This is the first time that these two polymorphisms have been found unlinked in nature, and the isolate shows hallmarks of recombination with Clade B (see below) suggesting a mechanism by which these polymorphism have become unlinked. This isolate was resistant to itraconazole, voriconazole and posaconazole (MIC ≥ 16 mg/L; 4 mg/L; 1 mg/L respectively) showing that the tandem repeat alone is sufficient to confer resistance in this genetic background. Intriguingly, genome analysis of 17 isolates with raised MICs to one of more of the azole test drugs (i.e. 16% of total resistant strains) failed to detect any known resistance mechanism or polymorphism with the *cyp51A* gene, indicating the presence of non-*cyp51A*-based azole resistance mechanisms among the UK and Ireland sample collection (Table 3). All of these isolates were obtained from clinical sources.

### Phylogenomics and signatures of directional selection associated with azole-resistance

Phylogenetic analysis showed a distinctive ‘dumbbell’ shape in the unrooted phylogeny similar to that reported by Abdolrasouli *et al.*^21^, and also identified two broadly divergent clades (Figures 1 and S2) with high (100%) bootstrap support. ‘Clade A’ contained 123 isolates, whereas ‘Clade B’ contained 95 isolates; Table 1 contains the breakdown of frequencies within both of these clades. The majority (99/123; 80%) of resistance-associated genotypes were clustered within Clade A of the phylogeny, in which 80% of isolates (*n* = 99/123) exhibited polymorphisms in *cyp51A* resulting in an azole resistant phenotype (Figures 1 and S2; Table 1). Conversely, the majority of isolates with no azole-associated resistance polymorphisms within *cyp51A* and an azole-susceptible phenotype clustered into the second clade, Clade B in which 86% of isolates (*n* = 82/95) exhibited an azole-susceptible phenotype.

**Figure 2.**
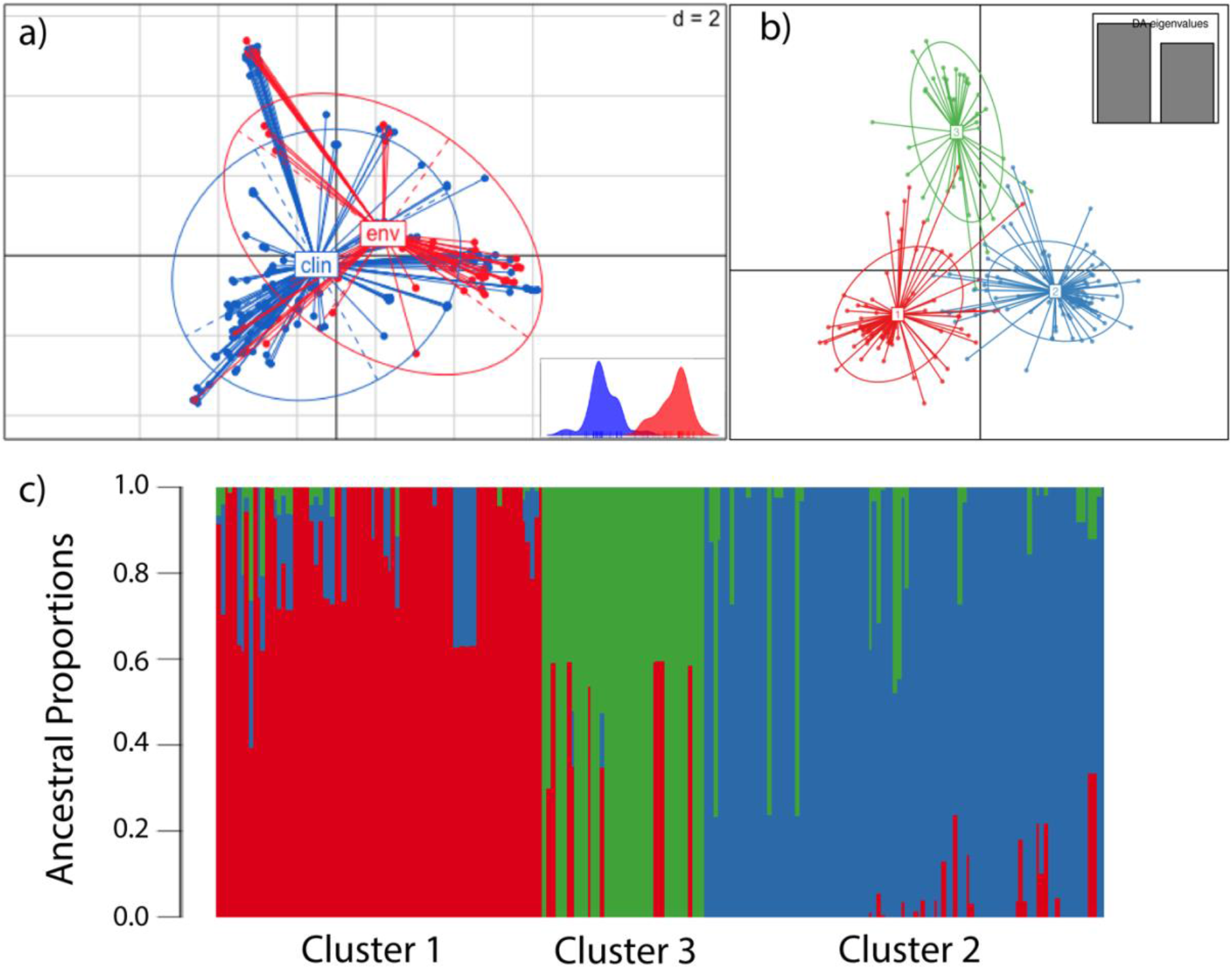
Occurrence of three subclusters within the *A. fumigatus* population, and clinical and environmental isolates are drawn from a single population. a) Scatterplot of the PCA of *A. fumigatus* genotypes using the first two principal components illustrating genetic identity for clinical and environmental isolates b) DAPC and PCA broadly identify three clusters, Clusters 1, 2 and 3, corresponding to the lowest Bayesian Information Criterion (BIC) c) Three subclusters are confirmed using STRUCTURE and *k* = 3

Whole genome SNP data were transformed using Principal Component Analysis (PCA) to identify the optimal number of clusters (*k*) corresponding to the lowest Bayesian Information Criterion (BIC). Here, we identified three clusters, containing a mix of environmental and clinical isolates in each, which broadly corresponded to the previous phylogeny (Figure S3): the first cluster corresponded to a subset of Clade A containing a broad selection of *cyp51A* polymorphisms (henceforth referred to as Cluster 1); the second cluster corresponded to Clade B (Cluster 2), and the third cluster overlapped with isolates within Clade A containing TR_34_/L98H only (henceforth referred to as Cluster 3). These genetic clusters displayed no geographical or temporal clustering (Figure S3). Discriminant Principal Component Analysis (DAPC) was used to examine the extent of divergence between genetic groups, whilst minimising variation within groups; this analysis confirmed the three genetically distinct clusters (Figure 2b). These three clusters were also confirmed using STRUCTURE and *k* = 3 (Figure 2c).

Minimum inhibitory concentrations (MICs) for the three test azole antifungal drugs, as well as the occurrence of polymorphisms associated with the c*yp51A* gene, were significantly higher in Clade A overall than in Clade B (*χ*^2^ test *p*-value = 3.27308e^-14^, df = 1). For tandem-repeat associated polymorphisms, 100% of the TR_34_ and 71% (*n* = 5) of the TR_46_-associated alleles occurred within Clade A, strikingly showing a complete absence of the commonly occurring TR_34_ genotype from Clade B).

We found that less than a quarter (23%; *n* = 75,317 SNPs) of the total *A. fumigatus* diversity seen in the whole dataset occurs within Clade A, despite this clade comprising 56% of the total isolates sampled. Of this nucleotide diversity within Clade A, no SNPs were uniquely associated with TR_34_/L98H or TR_46_ polymorphisms (the latter are distributed across Clades A and B). These data mirror those found by STR*Af* microsatellite analysis, illustrating that isolates harbouring drug resistance polymorphisms display reduced genetic diversity and are genetically depauperate in comparison to randomly-selected wild-type isolates^27^.

We estimated the index of association (I_A_) and the modified statistic rBarD^45^ in order to determine whether a signal of recombination is present within Clusters 1, 2 and 3. We assumed the null hypothesis of no linkage disequilibrium, meaning therefore that no recombination would not be rejected if the resulting values of both statistics were not significantly different from the distribution of values obtained from 999 samplings. For all three clusters, the null hypothesis could not be rejected, implying no significant linkage among the loci, and therefore that all three clusters are recombining with each other. In order to further explore recombination within *A. fumigatus* in the context of its population structure, we implemented a chromosome painting approach using *fineStructure*^43^, which identified shared genomic regions between individuals and populations. The linked coancestry model found the highest level of sharing within clades (Figure 3). Individual isolates C4, C54 and C178 within Clade B displayed evidence for strong haplotype sharing. These three isolates are all *MAT1-2* idiomorph. C4 was isolated from a soil sample in South Wales, C178 in a London hospital (a clinical isolate) and C54 is a clinical isolate from Dublin, Republic of Ireland (Supplementary Table 1). Overall, however, strong haplotype donation occurred mostly within clades; an exception to this observation was the strong haplotype donation between C178, C4, C54 (Clade B isolates) and C365 (a Clade A isolate). C365 is an environmental itraconazole resistant isolate (MIC ≥ 16 mg/L) with only TR_34_ present within *cyp51A*, and *MAT1-1* mating idiomorph.

**Figure 3.**
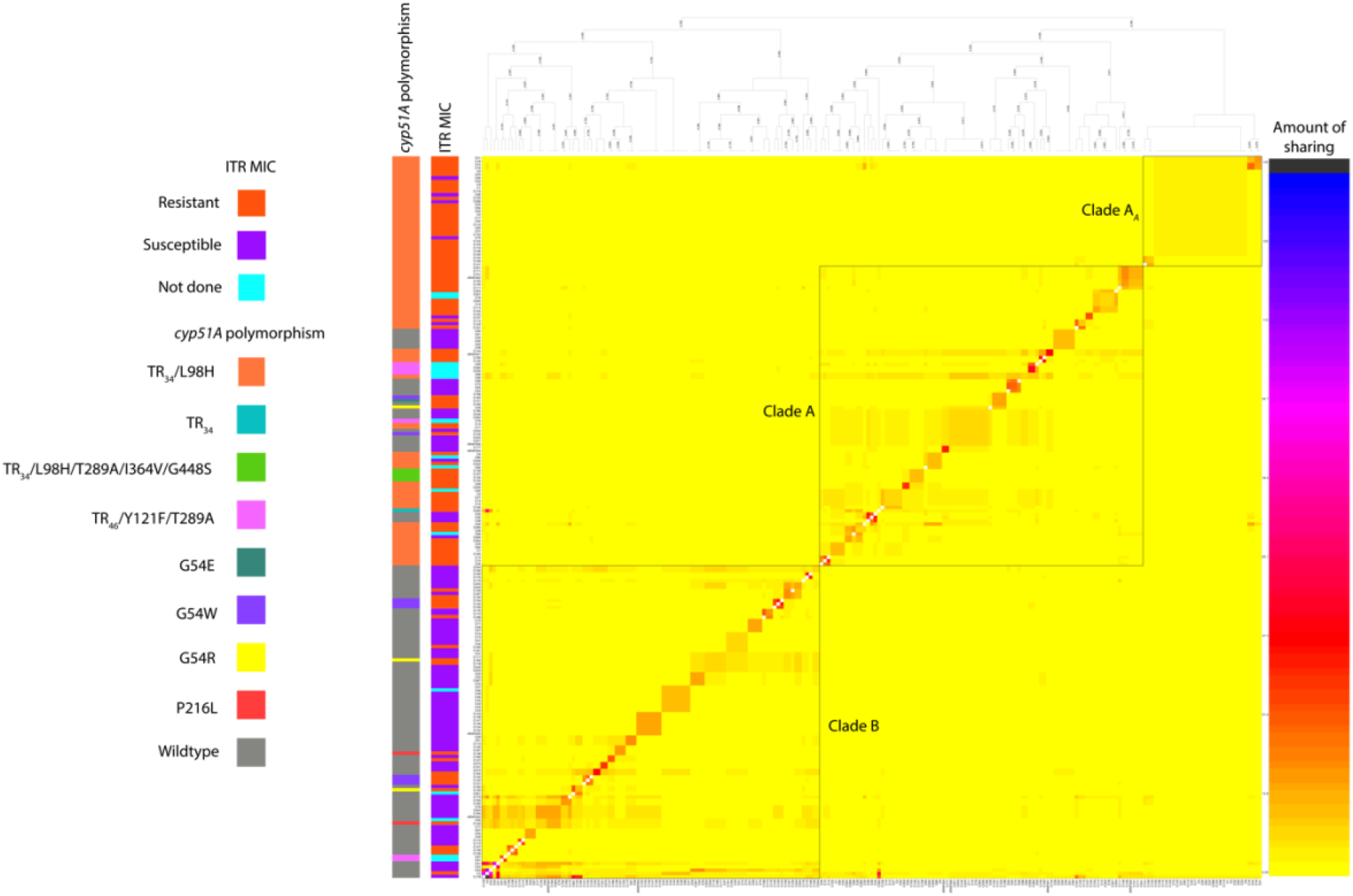
Genome sharing fineStructure analysis of *A. fumigatus* using genome-wide SNPs confirms the presence of three populations within the dataset. Population averaged coancestry matrix for the linked model dataset with associated *cyp51A* polymorphism and itraconazole MIC (defined as above or below 2 mg/L for resistance or susceptibility, respectively, or not done). The righthand scale bar represents the amount of genomic sharing, with blue/black representing the largest amount of sharing of genetic material, and yellow representing the least amount of shared genetic material.

We next performed further population-level genome-wide analyses in order to investigate whether local regions of the genome were differentially associated with respect to clade and resistance phenotype. We measured signatures of genome-wide population differentiation *via* the fixation index (F*_ST_*) analysis of non-overlapping 10 kb windows by comparing isolates within Clade A against those isolates from Clade B (Figure 4b). The average F*_ST_* value was 0.1312 (range: from 0 to 0.944035; standard deviation 0.0823); average F*_ST_* values and ranges for each chromosome are detailed in Table S3. Regions of extremely variable F*_ST_* values were observed in Chromosome I (range 0 to 0.9440). In particular a region of 590 kbp on the right arm of this chromosome displayed an average F*_ST_* value of 0.2273, but with a range in F*_ST_* values from 0 to 0.944035, suggesting near panmixis in some parts of this region between Clades A and B. Across this region 184 genes are found (gene ID Afu1g15860 to Afu1g17640 (Supplementary Data 4)). Of these genes, 9 contained SNPs which were found to be significantly associated with itraconazole resistance using treeWAS. Also within this region in Chromosome I, three extremely high outlier F*_ST_* values were observed where the average F*_ST_* value was 0.9321 (range: 0.9238-0.9440) with the spanned regions containing 9 genes (Table S4).

**Figure 4.**
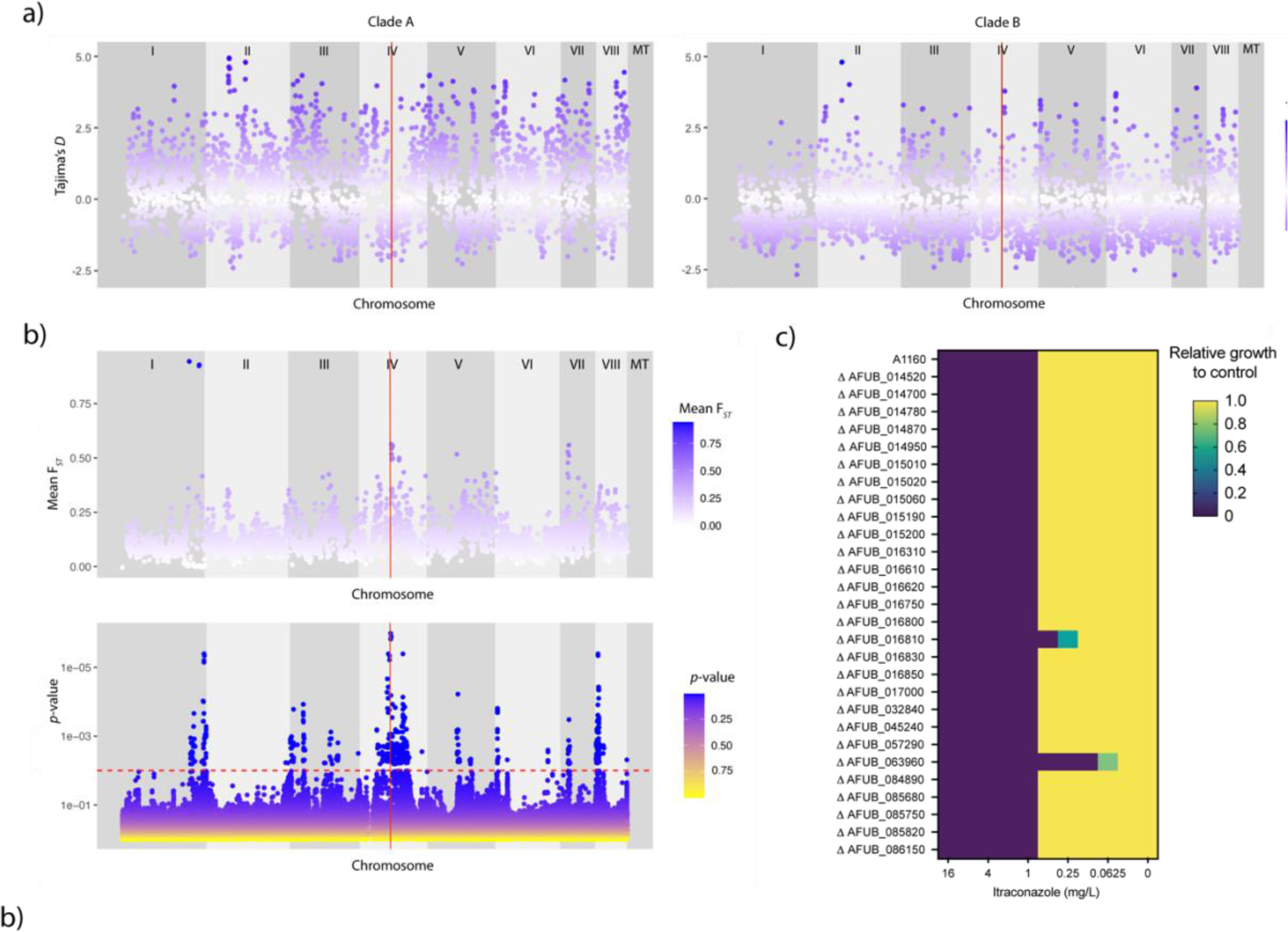
Loci associated with itraconazole resistance linked to regions of high FST and selection. a) Scatterplot of Tajima’s *D* estimates for each chromosome for all isolates within Clade A (left) and scatterplot of Tajima’s *D* estimates for each chromosome for all isolates within Clade B (right). The position of *cyp51A* is highlighted in red. b) Scatterplot of sliding 10-kb non-overlapping window estimates of F*_ST_* for each chromosome between isolates within Clade A and B (top panel). Manhattan plot (bottom panel) for *treeWAS* subsequent test (bottom panel) showing *p*-values for all loci, a significant threshold of 0.01 (dashed red line), above which points indicate significant associations. The vertical red line in both plots denotes the position of *cyp51A*. c) Relative growth of null mutants (compared to A1160) of genes with significant loci identified in *treeWAS* on media containing itraconazole.

**Table 4:** MIC that null mutants of genes identified as being statistically significant loci by treeWAS stop growing at, against control A1160

| A1160 gene ID | Af293 gene ID | MIC<br>(mg/L) | treeWAS *p*-<br>value | Gene description | Loci |
| --- | --- | --- | --- | --- | --- |
| AFUB_063960 | AFUA_4G06890 | >0.0625 | <2.03<sup>e-06</sup> | *cyp51A* | L98H |
| AFUB_016810 | AFUA_1G17440 | >0.25 | 0.001546976 | *abcA* | Y1149N |
| AFUB_014520 | AFUA_1G14960 | >1 | 0.001546976 | Has domain(s) with predicted actin binding activity | A820V |
| AFUB_014700 | AFUA_1G15150 | >1 | 0.003190639 | Ortholog(s) have hydrolase activity | L100F |
| AFUB_014780 | AFUA_1G15230 | >1 | 0.004462783 | Ortholog(s) have role in conidiophore development | Synonymous SNP |
| AFUB_014950 | AFUA_1G15410 | >1 | 0.000502209 | Has domain(s) with predicted zinc ion binding activity | S109P |
| AFUB_015010 | AFUA_1G15460 | >1 | 0.000200224 | Ortholog of *A. nidulans* FGSC A4: AN0892 | Synonymous SNP |
| AFUB_015020 | AFUA_1G15470 | >1 | 0.005395385 | Ortholog(s) have RNA polymerase II transcription factor activity | Synonymous SNP |
| AFUB_015060 | AFUA_1G15520 | >1 | 0.001956814 | Ortholog(s) have allophanate hydrolase activity | F2S |
| AFUB_015190 | AFUA_1G15660 | >1 | 0.005395385 | Ortholog of NRRL 181: NFIA_009670 | L71S |
| AFUB_015200 | AFUA_1G15670 | >1 | 0.001956814 | Putative laccase | Synonymous SNP |
| AFUB_016310 | AFUA_1G16920 | >1 | 0.005567187 | Beta-xylosidase | Synonymous SNP |
| AFUB_016610 | AFUA_1G17220 | >1 | 0.0000936 | Putative secreted polygalacturonase GH-28 | Intron |
| AFUB_016620 | AFUA_1G17230 | >1 | 0.000359083 | Ortholog(s) have carbon-oxygen lyase activity | P466T |
| AFUB_016750 | AFUA_1G17380 | >1 | 0.000200224 | Has domain(s) with predicted oxidoreductase activity | I68V |
| AFUB_016800 | AFUA_1G17430 | >1 | 0.00000228 | Ortholog(s) have monophenol monooxygenase activity | E586D |
| AFUB_016830 | AFUA_1G17470 | >1 | 0.00000305 | *nrtB* | Intron |
| AFUB_016850 | AFUA_1G17490 | >1 | 0.001152366 | Has domain(s) with predicted carbohydrate binding | G298S |
| AFUB_017000 | AFUA_1G17620 | >1 | 0.000887684 | Ortholog of RIB40: AO90011000197 | Synonymous SNP |
| AFUB_032840 | AFUA_2G17190 | >1 | 0.005886174 | Has domain(s) with predicted ATP binding | P1013A |
| AFUB_045240 | AFUA_3G03010 | >1 | 0.000887684 | Putative phosphate-repressible phosphate permease | Synonymous SNP |
| AFUB_057290 | AFUA_5G09740 | >1 | 0.004617329 | Ortholog(s) have role in conidiophore development | E432K |
| AFUB_084890 | AFUA_8G01700 | >1 | 0.003128212 | Has domain(s) with predicted catalytic activity | Synonymous SNP |
| AFUB_085680 | AFUA_8G00900 | >1 | 0.002175055 | Ortholog of FGSC A4: AN8368 | T47M |
| AFUB_085750 | AFUA_8G00820 | >1 | 0.000397402 | Ortholog of RIB40: AO90138000119 | H16R |
| AFUB_085820 | AFUA_8G00750 | >1 | 0.0000406 | Has domain(s) with predicted DNA binding | Synonymous SNP |
| AFUB_086150 | AFUA_8G00420 | >1 | 0.001260979 | C6 finger transcription factor *fumR* | N305D |

Regions of higher than average F*_ST_* (around 0.5) were also observed in Chromosomes IV and 7 (Figure 4b). The average F*_ST_* value for the *cyp51A* region, found in Chromosome IV, was 0.1193, slightly less than the average F*_ST_* value for the whole dataset (0.1312).

In order to investigate whether a signature of selection was observed across the genome of UK *A. fumigatus*, we used Tajima’s *D*^58^ statistic to measure departures from neutral expectations in non-overlapping 10 kb windows in order to measure evidence for selection when subsetting the dataset into Clade A and B populations. Across our dataset, we found highly variable positive and negative *D* values for both Clades (Figure 4a) that are suggestive of different patterns of demography and natural selection. Strikingly, on average, Tajima’s *D* was 0.4766 for Clade A and −0.3839 for Clade B, and observation was apparent per chromosome across Clades A and B (Table S3). The region in Chromosome IV containing *cyp51A* appeared to have a lower than average *D* value (−1.02123; Figure 4a) when compared against the rest of Clade A.

In order to determine the extent that variation in Tajima’s *D* and F*_ST_* owe to intra-clade population substructure, we subset the dataset into the three clusters that were defined by PCA, DAPC and STRUCTURE. As before, we used non-overlapping 10 kb sliding windows to estimate the two statistics. Three-way F*_ST_* (Figure S4) between the three clusters (Cluster 1 and Cluster 3, and Clade B (Cluster 2)) shows regions in Chromosome I that are approaching F*_ST_* of 1, showing that these highly-diverged alleles are present across Clade A. There were highly variable F*_ST_* values when comparing Clusters 1 and 3 (range: from 0 to 0.786203; average: 0.1309), Clusters 1 and 2 (range: 0 to 0.9322; average: 0.1577), and Clusters 2 and 3 (range: 0 to 1; average: 0.1424). Estimating Tajima’s *D* in Clusters 1, 2 and 3 also found highly variable positive and negative values of *D* across all three clusters (Figure S4). The average value of *D* for all three clades was around zero, a marked departure from the previous analysis of solely Clade A (Cluster 1: −0.089152, range −2.40694 to 4.68301; Cluster 3: 0.01773, range −2.62465 to 4.901) suggesting that the positive signature of selection in Clade A is largely owed to population substructure.

### Identifying loci significantly associated with itraconazole resistance

*treeWAS* is a microbe-specific approach that utilises a phylogeny-aware approach to performing genome-wide association whilst being robust to the confounding effects of clonality and genetic structure. We applied *treeWAS* to all sequenced isolates with reported itraconazole MICs and a binary phenotype (205 out of 218 isolates). The binary phenotype was categorised according to the itraconazole MIC defining susceptibility as MIC < 2 mg/L and resistant as MIC ≥ 2 mg/L. A phylogeny for these isolates was reconstructed as previously described with 281,874 SNPs common to one or more isolates with reported itraconazole MIC.

Analysis of this dataset led to the identification of 2,179 significant loci using the subsequent test; this test has been previously defined by Collin & Didelot^46^ as most effective at detecting subtle patterns of association. Sixty-four percent of significant loci (*n* = 1,391) were found to be intergenic. The three most significant loci (*p* < 2.03^e-06^) were all located on Chromosome IV: one was intergenic (position 1779747), one was a non-synonymous SNP (S237F) in Afu4g07010 (position 1816171) and the third was the L98H substitution in *cyp51A* (position 1784968 on chromosome IV; Supplementary Data 1). Of the significant loci identified on other chromosomes, some mapped to genes involved in secondary metabolism, including fumitremorgin. There were also loci within Afu8g00230, encoding verruculogen synthase, which is associated with *A. fumigatus* hyphae and conidia modifying the properties of human nasal epithelial cells^59^. A single significant SNP was also identified in *Aspf2* (Afu4g09580), a gene which is involved in immune evasion and cell damage^60^.

A striking pattern was observed whereby peaks of gene-phenotype associations on Chromosomes I, IV and VII mirrored regions of high F*_ST_* when comparing Clades A and B (Figure 4b) likely reflecting the impact of azole selection upon these alleles. Of these regions, 1,385 out of the 2,179 significant loci were located within Chromosome 4. An exception to this was observed in Chromosome VIII, where F*_ST_* values did not peak above 0.370951, but *treeWAS p*-values were significant. Significant loci within Chromosome VIII are located in genes such as verruculogen synthase, PKS-NRPS synthetase *psoA* (Afu8g00540) and brevianamide F prenyltransferase (Afu8g00210). Out of the 356 significant loci in Chromosome VIII, the majority (*n* = 354) were located within 500,000 bp of the start of the chromosome, and 62% were intergenic.

In order to determine whether there was an observable link between the significant loci identified by *treeWAS* and azole-resistance, preliminary investigations were undertaken using 28 gene deletion mutants from the COFUN knockout collection in which the corresponding loci were deleted. These mutants were screened for growth on media containing itraconazole relative to the parental control (strain A1160). As expected, null mutant ΔAFUB_063960 (*cyp51A*) was not able to fully grow in media containing >0.06 mg/L of itraconazole; in addition null mutant ΔAFUB_016810 (*abcA*) was unable to grow in media containing >0.25 mg/L of itraconazole (Figure 4c). The control strain and all other null mutants were unable to grow in media containing >1 mg/L (Figure 4c, Table 4).

### Clinical and environmental *A. fumigatus* consist of genotypes with high relatedness

We subset our dataset into genotypes from clinical and environmental sources, then applied PCA and DAPC. Both metrics showed a lack of genetic differentiation amongst isolates from clinical and environmental origins showing, importantly, that clinical isolates are drawn from the wider environmental population (Figure 2a). On average, any pair of isolates within the dataset were separated by 11,828 SNPs. Within the phylogeny, out of 218 isolates, 6 pairs or groups of *A. fumigatus* contained both clinical and environmental isolates that were genetically very highly related (Table S5), with bootstrap support of 65% or higher (range: 65-100%: median: 100%), from both Clades A and B. For these paired or grouped environmental and clinical isolates, average pairwise diversity was 297 SNPs (compared to an average pairwise diversity of 11,828 SNPs observed for the complete dataset). Out of these highly-related pairs and groups that share both clinical and environmental isolates, four contained TR_34_/L98H. On average, any clinical/environmental pair or group of the azole-resistant isolates were separated by 247 SNPs (range: 227-270 SNPs; standard deviation: 19).

The pair or group showing the highest identity were Group 5 in Clade B, containing isolates C42, 43, 44, 48 (isolated from a CF patient in Dublin, Republic of Ireland) and 96 (isolated from a plant bulb in Dublin, Republic of Ireland). The isolates within Group 5 were separated by only 217 SNPs (1.8% of the total diversity seen in this dataset), were all *MAT1-2* mating type, and did not contain any polymorphisms within *cyp51A* associated with drug resistance, nor had raised MICs (Table S2).

We discovered a further cluster of very highly related *A. fumigatus* isolates consisting of 14 clinical and 14 environmental isolates that all harboured the TR_34_/L98H c*yp51A* polymorphism. These isolates were also all *MAT1-2* mating type. This clonal clade sits within the larger Clade A and within Cluster 3, and will henceforth be referred to as Clade A*_A_* (Figure 1). Isolates within Clade A*_A_* were found to be broadly distributed across England, Wales, Scotland and the Republic of Ireland (Figure 1b), covering a spatial distance equivalent to that observed across the whole dataset (Figure 1b). The clonal Clade A*_A_* cluster appears with high frequency, comprising 13% of the total dataset and 23% of the isolates found within Clade A. Clade A*_A_* isolates display low genetic diversity and high bootstrap support (100%) within the phylogeny; The average number of SNPs separating Clade A*_A_* isolates is 294 SNPs and contains 0.09% of total genetic diversity observed in the whole dataset. Isolation by Distance (IBD) was examined *via* a MANTEL test implemented in adegenet^41^ in R, and showed no significant correlation (Observation = 0.0379 with a simulated *p*-value of 0.01) between genetic and geographical distances suggesting that the Clade A*_A_* clone will have a broader extent than just the British Isles.

Clade A*_A_* is significantly overrepresented (*χ*^2^ test *p*-value = 2.75e^-13^, df = 1) for drug-resistant environmental isolates, as all isolates contain TR_34_/L98H, compared to only 63% of environmental isolates in the rest of the dataset. In comparison, only 54% of clinical isolates within the dataset as a whole contain the TR_34_/L98H polymorphism. There were two instances within Clade A*_A_* where pairs of isolates from clinical and environmental sources showed high genetic identity between clinical and environmental isolates, Pairs 1 and 2. Of these two pairs, Pair 1 showed the highest identity, with just 227 SNPs separating environmental isolate C21 (isolated from soil in a potato field in Pembrokeshire, Wales) and clinical isolate C112 (isolated from a CF patient and supplied by the Public Health England National Mycology Reference Laboratory – location of original patient unknown). Previous studies have used nucleotide diversity (π) as a metric to test whether pairs of isolates are epidemiologically linked in order to infer transmission^61, 62^. Here, nucleotide diversity (π) tests implemented in vcftools^44^ showed that the genetic diversity separating the isolates in Pair 1 were significant different to the mean nucleotide diversity within the rest of the isolates within Clade A*_A_* (one-tailed t-test *p* < 3.0473^e-252^, Table S5), showing that these isolates are genealogically tightly linked and that this shared identity could not have occurred by chance occurrence alone.

## Discussion

Emerging resistance to antifungal drugs is compromising our ability to prevent and treat fungal diseases^7^. Breakthrough infections by azole-resistant *A. fumigatus* phenotypes have been observed with striking increases across northern Europe where the incidence has increased from negligible levels pre-1999 up to measured prevalences of 3-40% in the present day^11, 63^. Further, the advent of COVID-19 has created a large and growing global cohort of patients that are at risk of azole-resistant *A. fumigatus* coinfections^64^. Our UK-wide genomic analysis of environmentally- and clinically-sourced *A. fumigatus* yielded two main findings. Firstly, *A. fumigatus* showed strong genetic structuring into two Clades ‘A’ and ‘B’, with the majority of environmentally-occurring azole-resistance alleles segregating inside Clade A and showing signatures of selection at multiple loci, some of which are known to adapt in response to selection by fungicides. Secondly, we observed multiple exemplars of drug-resistant genotypes in the clinic that matched those in the environment with very high identity. As patients have never convincingly been shown to transmit their *A. fumigatus* to the environment, this finding demonstrates that at-risk patients were infected by isolates that have pre-acquired their resistance to azoles in the environment.

Phylogenomic analysis confirmed that the population of *A. fumigatus* was not panmictic, and is structured into a characteristic ‘dumbbell’ phylogeny marked by two clades with limited inter-clade recombination. We used four hypothesis-free population genetic methods, PCA, DAPC, STRUCTURE and *fineStructure* to independently confirm the existence of the two clades, as well as the occurrence of strong genetic subclustering within Clade A and evidence of widely occurring clonal genotypes (*eg.* Clade A*_A_*). Azole resistance was defined both genetically, by the occurrence of known resistance alleles, as well as phenotypically using MICs and mapped onto the phylogenetic structure. This analysis showed that Clade A contained the majority of isolates (88%) harbouring polymorphisms in *cyp51A* associated with drug resistance compared to Clade B, which was predominantly wildtype for *cyp51A*. Phenotypically-defined resistance recovered a similar pattern with 78% of itraconazole MICs above clinical breakpoints being compartmentalised into Clade A. Further analysis using PCA, DAPC and STRUCTURE showed Clade A is divided into two subclusters, Clusters 1 and 3. All TR_34_/L98H polymorphisms were found within Cluster 3 while Cluster 1 contained isolates with a variety of *cyp51A* polymorphisms. Interestingly, isolates containing the TR_46_/Y121F/T289A polymorphism were found in both Clade B and Clade A Cluster 1, but not Clade A Cluster 3. Similarly, the novel polymorphisms TR_34_/L98H/T289A/I364V/G448S and the unlinked TR_34_ were found only in Clade A Cluster 3. These analyses suggest that TR-associated azole resistance has evolved a limited number of times and recombination has not yet had the impact of homogenising these alleles across the wider *A. fumigatus* phylogeny. Similar conclusions were drawn by Camps *et al.* based on microsatallite and CSP marker analysis of European isolates, who also suggested that the TR-resistance form had developed from a common ancestor or restricted set of genetically related isolates^24^. However, the allocation of these novel polymorphisms could also be due to not sampling the entire ecological space available. Future work on global collections of *A. fumigatus* that have been collected across longer periods of time than in the current study will however be needed to trace and date the spatiotemporal origins of these alleles.

Our dataset revealed new insights into the emerging complexity of *cyp51A-*encoded resistance polymorphism. By characterising the genotypes of *cyp51A* for all isolates we identified a novel *cyp51A* polymorphism TR_34_/L98H/T289A/I364V/G448S in four clinical isolates, which appears to be the result of recombination between genotypes containing the TR_34_/L98H, TR_46_/Y121F/T289A, and G448S polymorphisms. This polymorphism confers a pan-azole resistant phenotype, and is similar to the TR_46_ /Y121F/M172I/T289A/G448S polymorphism recently discovered in clinical isolates in the Netherlands^18^. Although this genotype has not yet been discovered in the environment, the occurrence of these alleles separately in environmental isolates suggests that it could have formed as a consequence of meiosis, and may be recovered given further surveillance. The counterargument, that this polymorphism evolved *de novo* in patient as a consequence of recombination *in vivo* is not impossible, but is considered rare. Such *de novo* evolution has been observed at other regions of the *A. fumigatus* genome^65^. We further identified, for the first time, a single environmental isolate in Scotland containing the TR_34_ tandem repeat by itself in *cyp51A*. This significant finding demonstrates that the tandem repeat in itself can confer azole-resistance as shown by its resistance to itraconazole (MIC 16 mg/L; Table S2 – isolate C365). It also suggests that either de-linkage from the L98H amino acid substitution occurs through recombination (in this case with the Clade B genetic background) or that parallel origin of the tandem repeat can occur through chance processes. While 86% of resistance to itraconazole defined using MICs was associated with known resistance alleles, we observed 12 cases where isolates were resistant to itraconazole yet without any known *cyp51A* polymorphisms associated with drug resistance (Table 3). Six of these isolates were also resistant to voriconazole and posaconazole, with two additional isolates displaying resistance to voriconazole and posaconazole only (Table 3). This finding, together with the identification of the TR_34_/L98H/T289A/I364V/G448S polymorphism, shows that we have not yet characterised the full spectrum of azole-resistance in *A. fumigatus* occurring in the environment posing a risk to the susceptible patient populations^18, 66^.

Utilising the chromosome painting approach *fineStructure* enabled the confirmation of three subpopulations within this dataset, which corresponded to Clades A, B and A*_A_*, and the presence of a subtle population substructure. Strong donation of haplotypes between Clade B isolates C4, C54 and C178 and the Clade A TR_34_-only isolate C365 is indicative of recent recombination between these two clades, highlighting the potential for recombination-driven resistance alleles to manifest. Previous studies have confirmed that asexual reproduction facilitates the emergence of TR_34_/L98H within *A. fumigatus*^67^, so it is likely this mechanism is also facilitating the emergence of new resistance alleles across the population, such as the novel alleles observed in this study. However, it remains to be seen whether sexual recombination is a mechanism responsible for facilitating the emergence and/or spread of resistance alleles within their clades. The lack of a reliable molecular clock for *A. fumigatus*, combined with evidence of recombination between the clades, currently hinders the dating of the time of emergence of resistance alleles.

Previous studies in bacteria, fungi and other eukaryotes have shown the advantages of using whole genome sequencing for investigating the effects of selection^68–70^. Calculating the population differentiation index *F*_ST_ on non-overlapping sliding windows between populations using genome-wide SNPs can detect regions of the genome that have been subject to stabilising or diversifying selection^71^, the impact of which can ultimately lead to the divergence of populations if recombination is rare. Our genome-scans show that Clade A and B explain ∼13% (*F*_ST_ = 0.13) of the observed genetic structure (per chromosome range 10% - 16%). This shows that, while the dumbbell phylogeny is not due to the presence of two sister and cryptic species (which would be marked by levels of differentiation approaching *F*_ST_ ∼ 1) the barrier to gene-flow is strong enough to allow these populations to experience the effects of selection differently. This is demonstrated by Tajima’s *D*, which switches sign between the two clades from *D =* 0.4766 for Clade A to *D =* −0.3839 for Clade B. The positive value for Clade A suggests an excess of intermediate-frequency SNPs, a distribution that is expected under a model of multiple clonal expansions of fit variants that are under directional selection. However, an excess of intermediate-frequency SNPs could also be explained by a mixture of populations; PCA, DAPC and STRUCTURE confirmed the presence of two clusters within Clade A; Cluster 1 which contains a variety of *cyp51A* polymorphisms, and Cluster 3, which contains isolates with the TR_34_/L98H polymorphism only. The positive value of Tajima’s *D* within clade A is largely dependent on the internal TR_34_/L98H – associated population structure and reduces to ∼0 when the clusters are analysed independently (Cluster 1 *D*: −0.089152, Cluster 3 *D*: 0.01773). Accordingly, it appears that the effects of fungicide selection upon the genetic background that these resistance alleles are found, combined with low rates of recombination, is driving much of the observed population genetic structure.

We challenged this observation by using a GWAS approach using the recently-developed microbe-specific tool *treeWAS*. This method maintains high statistical power while being robust to the confounding effects of clonality, population structure and recombination, which we find in this population of *A. fumigatus*. We found a striking congruence where significant peaks of gene-phenotype associations on Chromosomes I, IV and VII mirrored regions of high F*_ST_* when compared to Clades A and B (Figure 5a). This finding therefore links the phenotypes that we have measured (itraconazole breakpoints) to the regions of the genome that are under selection (chromosome IV *cyp51A* and other regions) and to the genetic structure that we observe (islands of divergence between Clade A and B containing significance *treeWAS* subsequent scores). Accordingly, it appears that fungicide-associated resistance is driving the patterns of evolution and is the most likely explanation for much of the observed genetic architecture.

Yet, there remains much to understand about how these genomic regions of divergence and signatures of selection govern azole resistance in chromosomes with no immediately obvious resistance functions. For instance *treeWAS* also identified significant SNPs in genes involved in secondary metabolism. Recent research has shown that secondary metabolites combat the host immune system and aid growth in the host (human) environment^72^. Our results identified four SNPs in the gene encoding fumitremorgin C monooxygenase (Afu8g00240), which part of the pathway for secondary metabolite fumitremorgin C, a mycotoxin that acts as a potent ABCG2/BCRP inhibitor that reverses multidrug resistance^73^. This pathway has also been suggested as regulating the brevianamide F gene cluster^74^; two non-synonymous SNPs in the gene encoding brevianamide F prenyltransferase (Afu8g00210) were also identified as significant in our *treeWAS* analysis. Further complexity is observed in Chromosome VIII, where F*_ST_* values did not peak above 0.370951, but *treeWAS p*-values were highly significant. Significant loci within Chromosome VIII are located in genes such as verruculogen synthase, PKS-NRPS synthetase psoA (Afu8g00540) and brevianamide F prenyltransferase (Afu8g00210). Out of the 356 significant loci in Chromosome VIII, the majority (*n* = 354) were located within 500,000 bp of the start of the chromosome, and 62% were intergenic. Relative growth of Δ*cyp51A* and Δ*abcA* null mutants were effected even at >0.06 mg/L and >0.25 mg/L respectively showing that these two genes have a key role in azole resistance and directly confirming the utility of *treeWAS* to identify loci associated with itraconazole resistance. Future work now needs to widen our search and to focus on the use of reverse functional genomic approaches to interrogate the function of these genes within the context of their potentially epistatic inter-relationships with the canonical ergosterol biosynthesis *cyp51A* azole-resistance alleles on Chromosome IV.

The ability of fungal pathogens to dispense with sex and to reproduce clonally through mitosis is an important biological feature that enables genotypes of high fitness to rapidly colonise new environments^75^. Examples include the invasion of North America by the fungus *Pseudogymnoascus destructans* causing bat white-nose syndrome^76^, *Cryptococcus gattii* invading the Pacific North West^77^, and panzootic amphibian chytridiomycosis globally. We identified a highly supported clade of 28 isolates of *A. fumigatus* demonstrating low genetic diversity within Clade A, Clade A*_A_*, which were all azole-resistant. These isolates were spatially unstructured and recovered from an area covering 63,497 miles^2^ across England, Wales, Scotland and Republic of Ireland, confirming finding from previous studies that azole-resistant *A. fumigatus* is capable of dispersing over wide geographical regions, perhaps even at a global level^21, 27^. That clinical and environmental isolates within Clade A*_A_* are genetically depauperate not only shows the impact of the genetic sweep that has accompanied the selection of these beneficial mutations, but also suggests that these isolates are highly fit in both clinical and environmental settings. These data indirectly corroborate previous findings that there are no obvious costs associated with the TR-related polymorphisms that might impact the virulence of these isolates^78, 79^.

Importantly, genetic similarity of paired isolates recovered from environment and clinical sources shows that acquisition of both wildtype and azole-resistant *A. fumigatus* has occurred widely across this dataset. The statistically significant association between the azole resistant isolates C354, C355 and C357, two clinical isolates and an environmental isolate respectively, isolated within the same city, coupled with their very high genetic similarity suggests an environment-to-patient acquisition of azole resistant *A. fumigatus* with very high confidence. The low number of SNPs separating the isolates, coupled with nucleotide diversity tests showed that these isolates are genealogically tightly linked, and were strongly supported phylogenetically with 100% bootstrap support also. This shared identity could not have occurred by chance alone, as this shared identity was found to be statistically significant (*p* < 3.0473^e-252^). Our PCA analysis showed that clinical isolates are drawn from a wider environmental diversity, with a lack of genetic differentiation amongst isolates in these populations (Figure 2a). The presence of other significant shared identity between clinical and environmental isolates (Table S5) demonstrates that this environmental to clinical acquisition *via* inhalation of fungal spores is not a rare occurrence.

Our study supports the hypothesis that the widespread use of azoles as fungicides in agriculture is coupled to the increasing isolation of azole-resistant *A. fumigatus* from environmental sources^14^. That these isolates bear hallmark multi-locus genotypes that are indistinguishable to those recovered from patients supports our conclusion that adaptation to fungicides in the environment is leading to acquisition of *A. fumigatus* bearing azole-resistance genotypes^26, 80^. Here, we also identify spatially widespread clones of *A. fumigatus* that are not only resistant to azoles but are also highly represented in both the environment and the clinic, suggesting that that there are few fitness costs associated with this phenotype. Respiratory viruses such as H1N1 influenza and Severe Acute Respiratory Syndrome (SARS) are known to predispose critically-ill patients to secondary mould infections^2^, and early reports suggest that similar infections will be experienced by patients with COVID-19^64^. The growing numbers of susceptible individuals underscores the need for further surveillance, which is acutely needed to more fully understand the risk posed by environmental reservoirs of pathogenic fungi that, through the use of agricultural antifungals, have evolved resistance to first-line clinical azoles.

## Acknowledgments

This study was partially supported by an unrestricted education grant from Gilead Science Ltd. through their investigator sponsored research programme. JR, TS, APB, PSD, DAJ and MCF were supported by a grants from Natural Environmental Research Council (NERC; NE/P001165/1 and NE/P000916/1), the UK Medical Research Council (MRC; MR/R015600/1) and the Wellcome Trust (219551/Z/19/Z). MCF is a CIFAR Fellow in the ‘Fungal Kingdom’ programme. KD was supported by a PhD studentship awarded by the School of Medicine, Trinity College Dublin. PGM and JR (Dublin) received a project grant from the National Children’s Hospital, Tallaght University Hospital, which in part supported this work. AW and EB are supported by the Wellcome Trust Strategic Award (grant 097377), the MRC Centre for Medical Mycology (grant MR/N006364/2) at the University of Exeter, and the BBSRC EASTBIO grant (BB/M010996/1).

## Supplementary Information

### Supplementary Tables

**Table S1.**
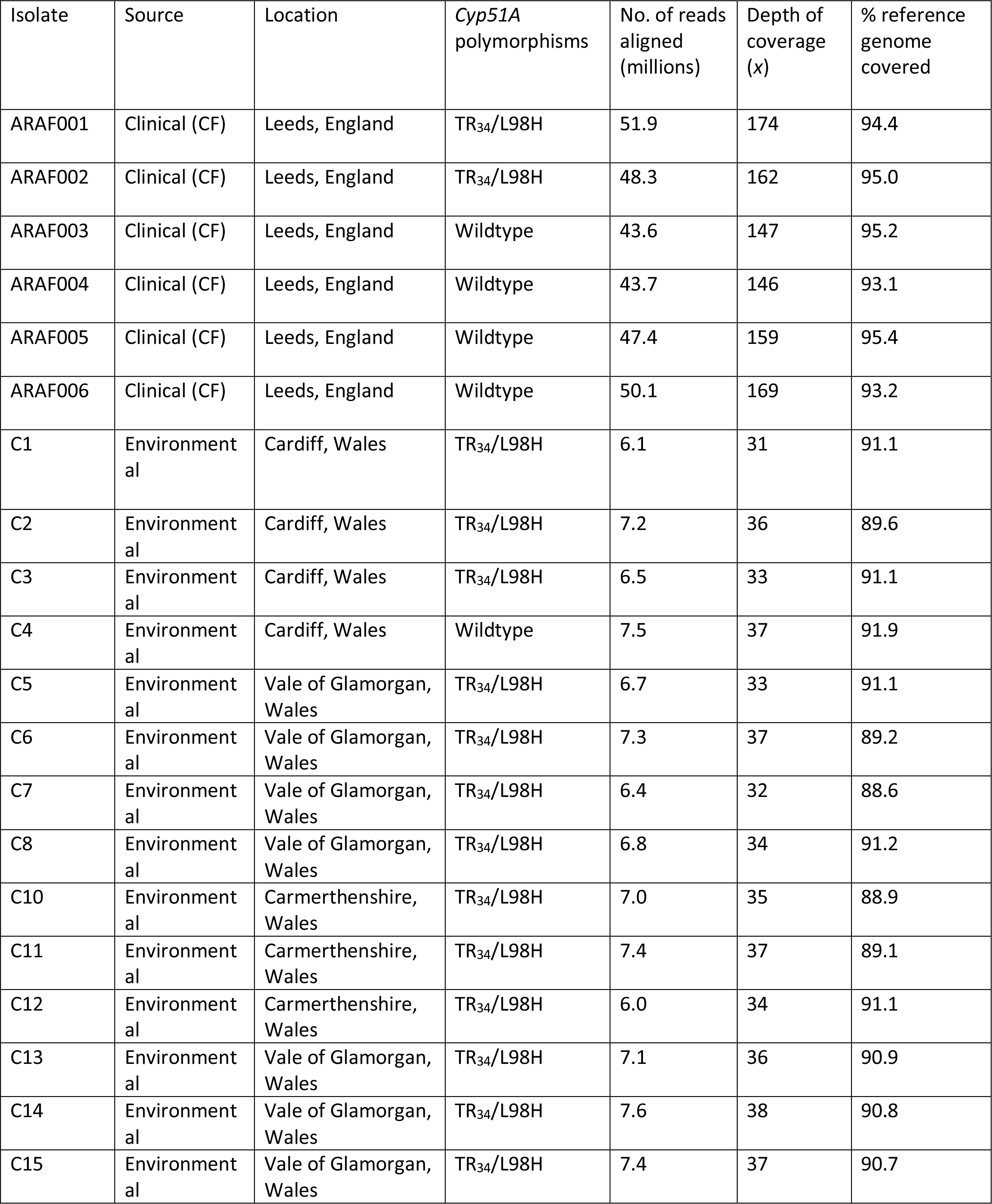
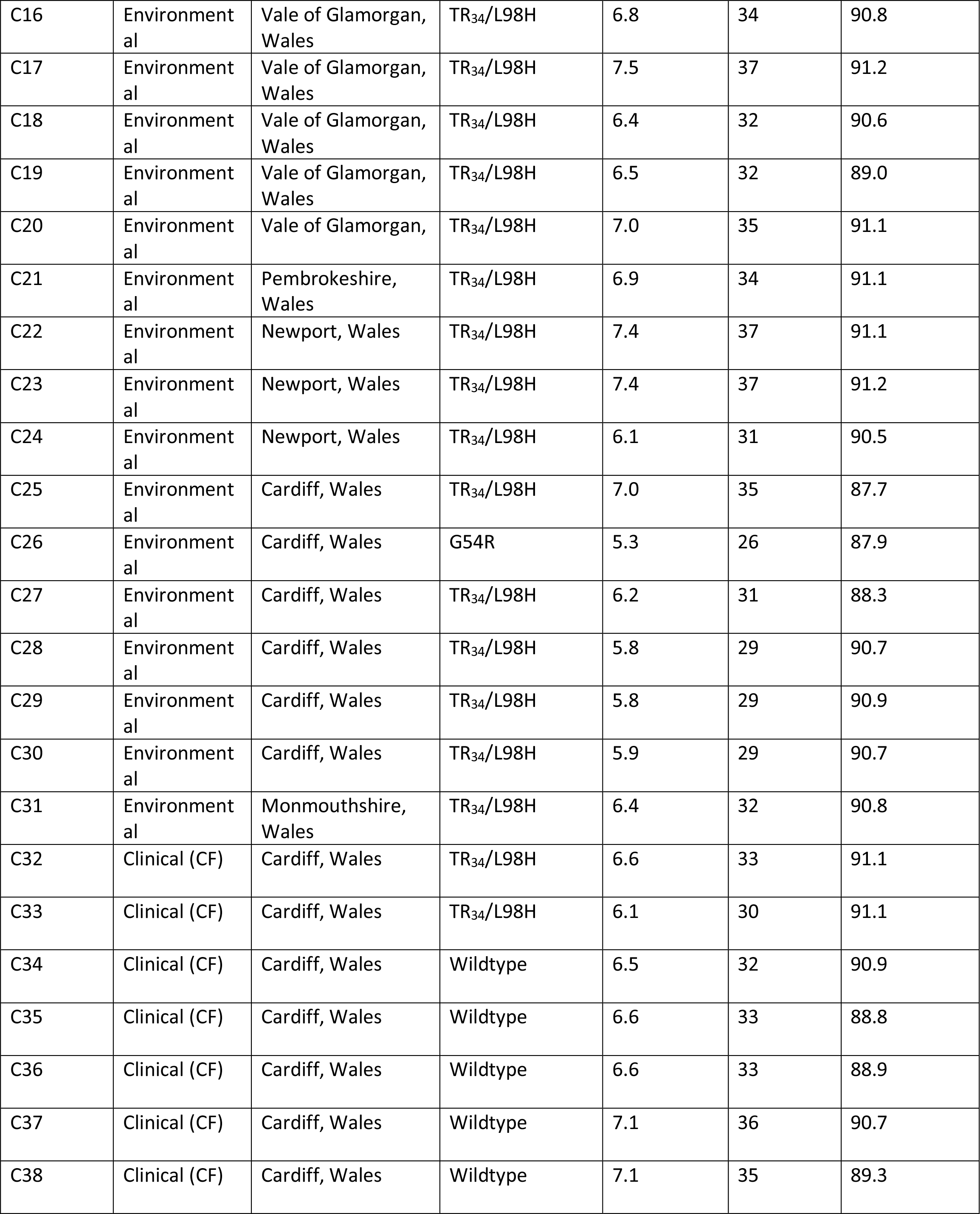
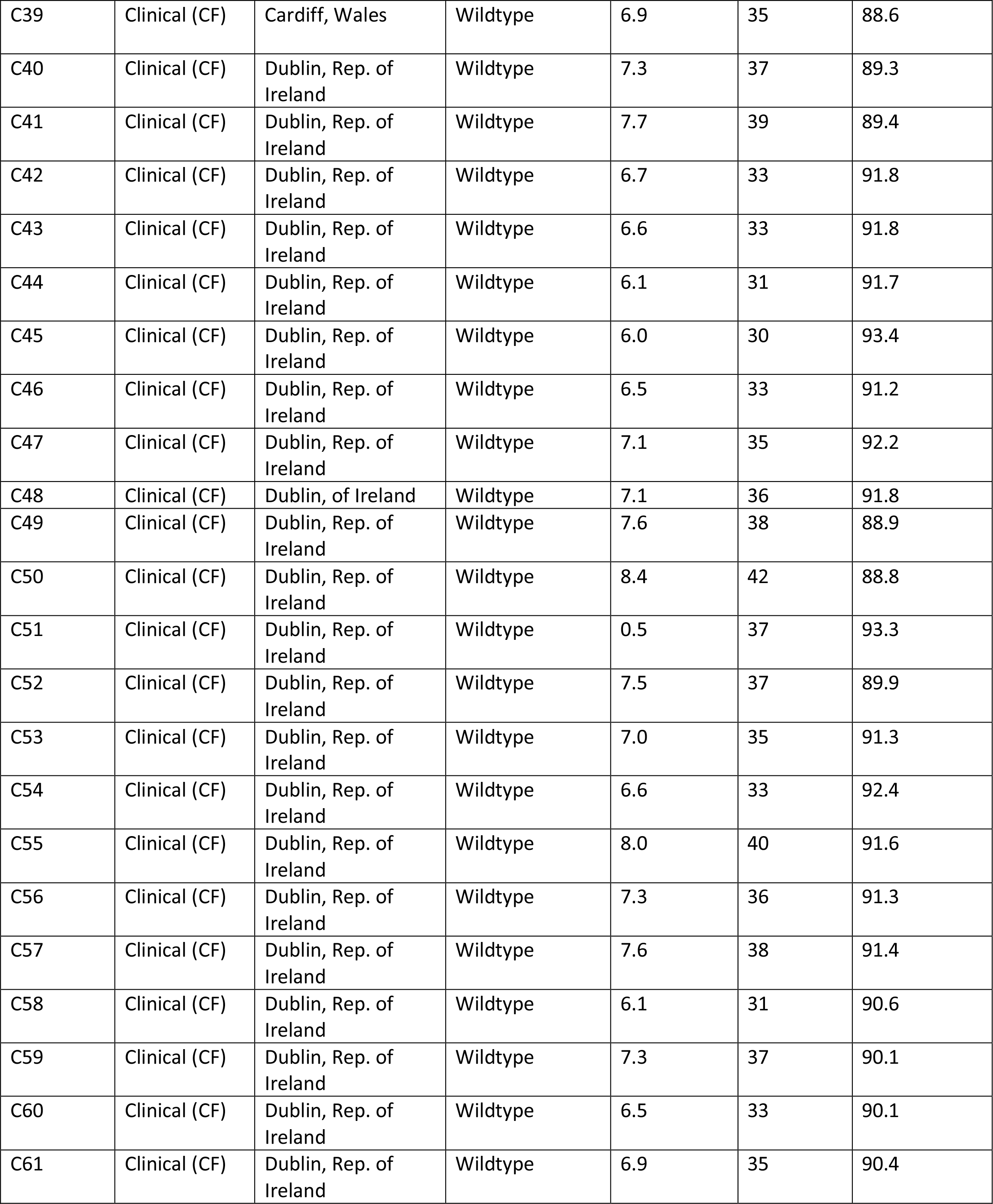
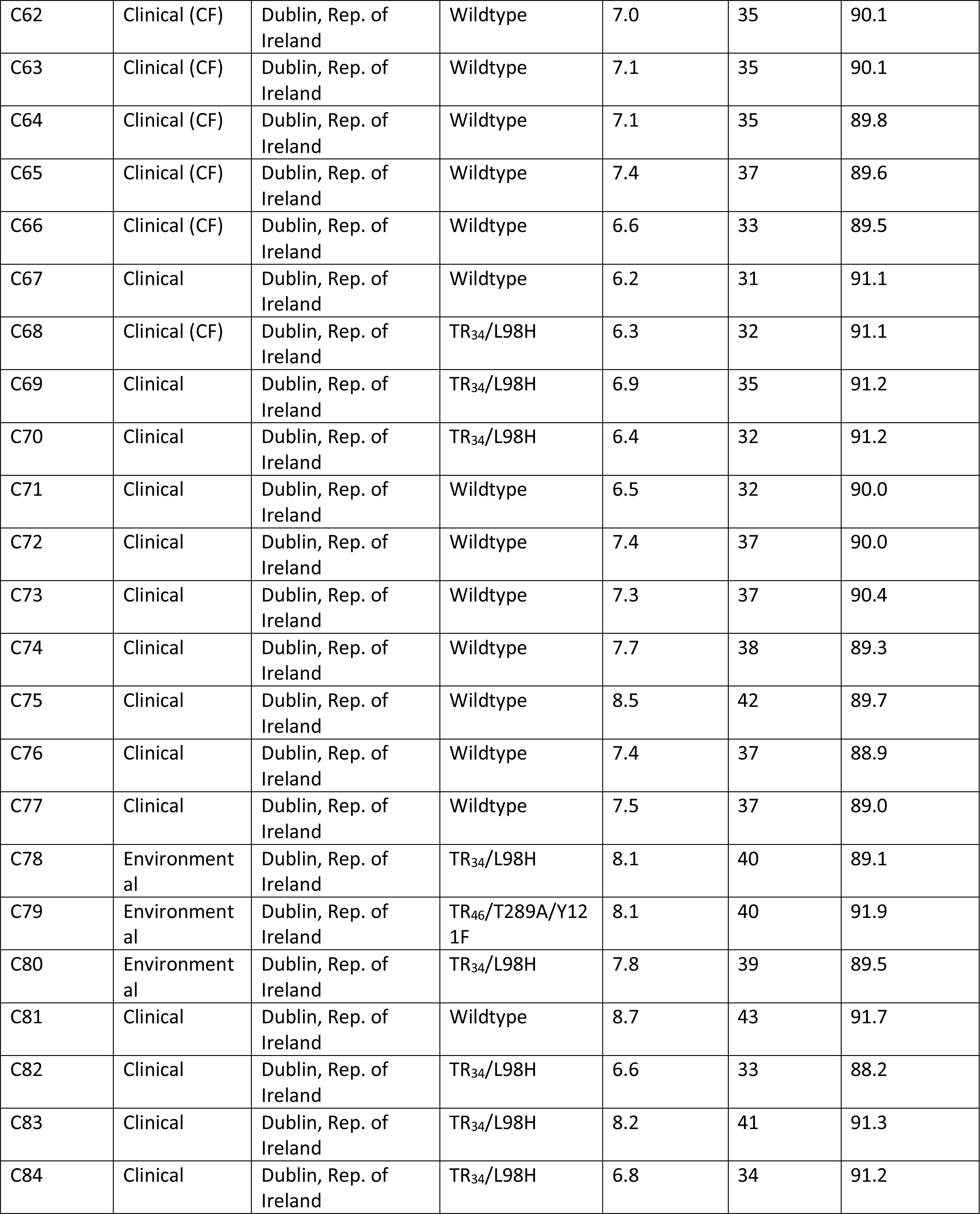
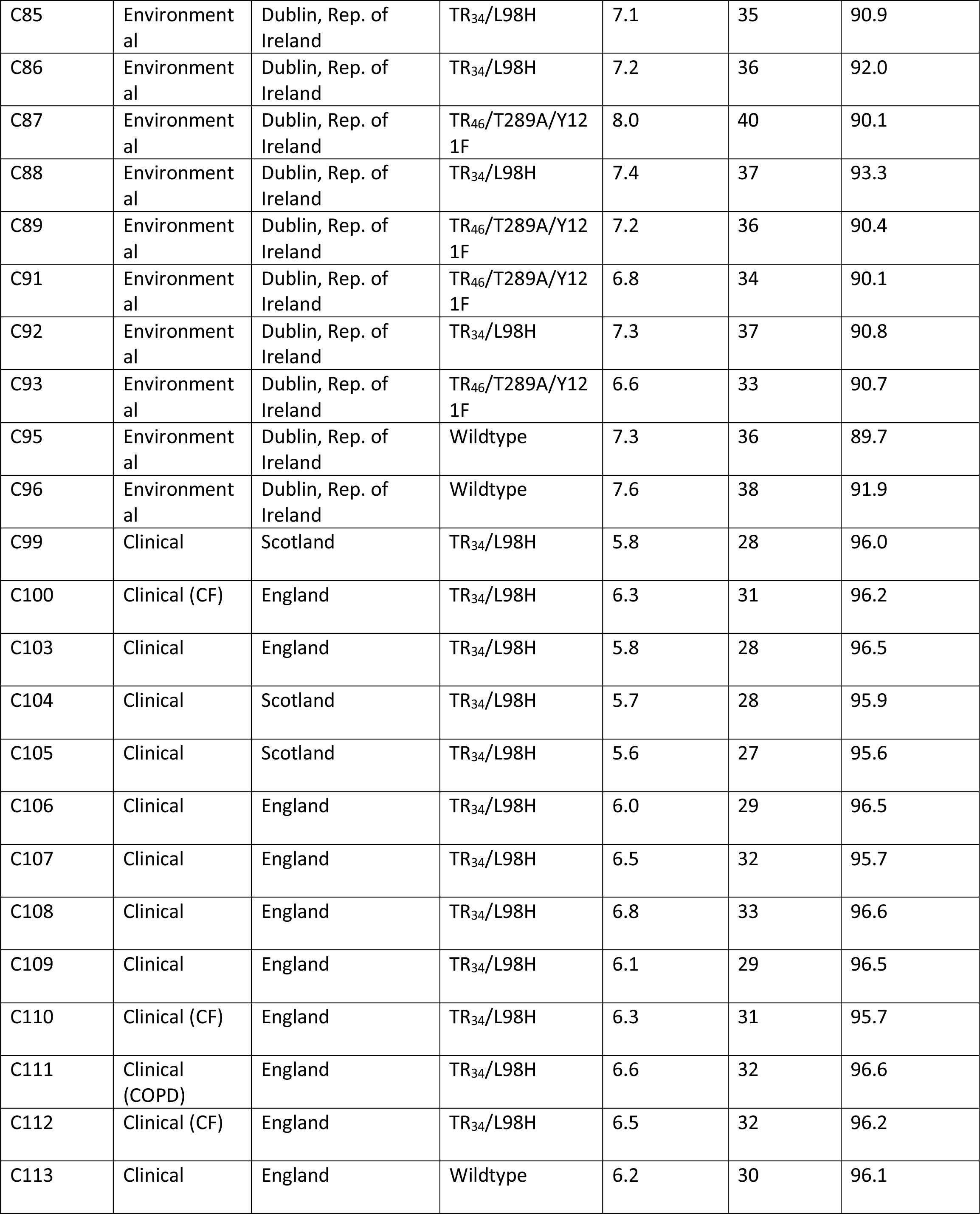
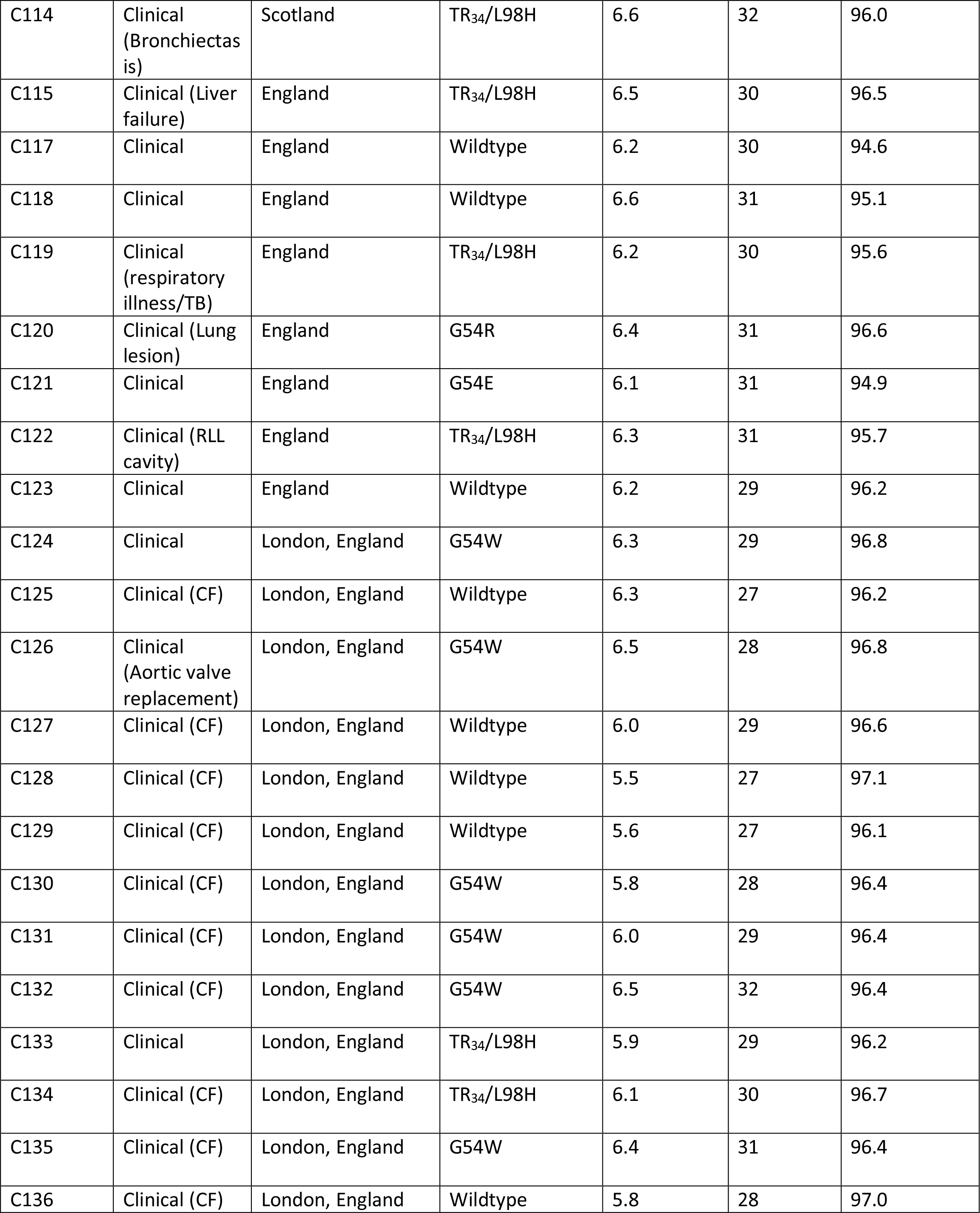
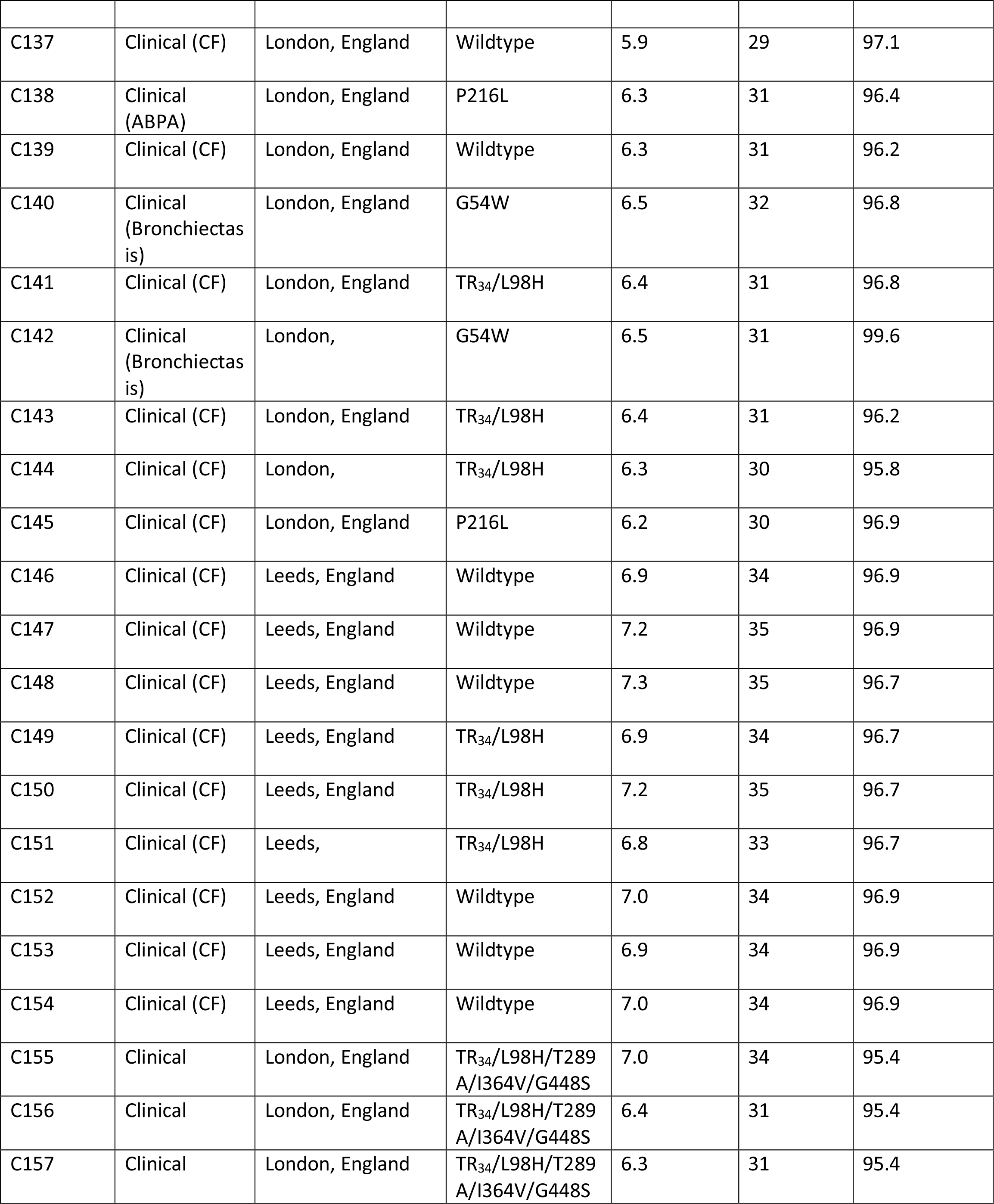
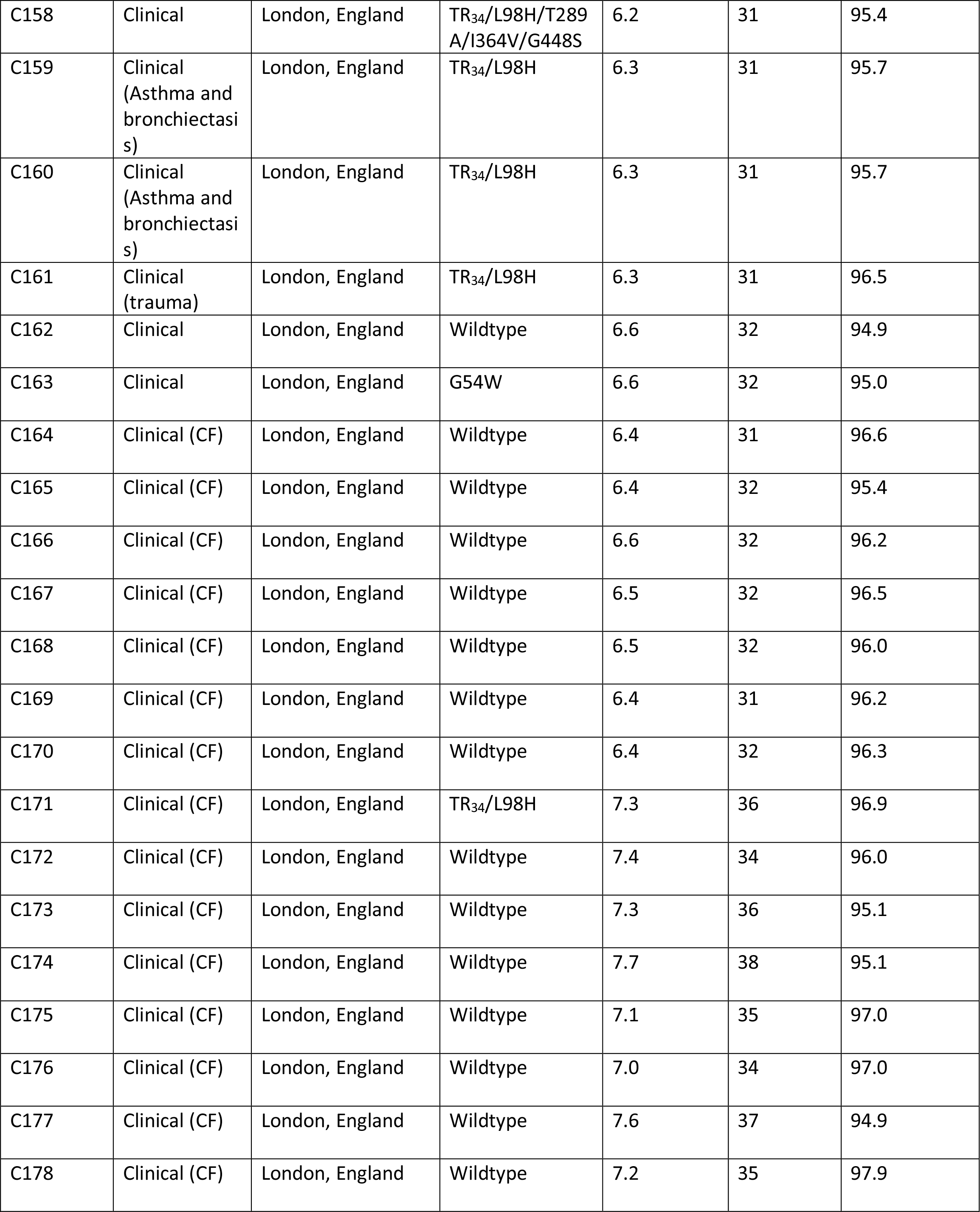
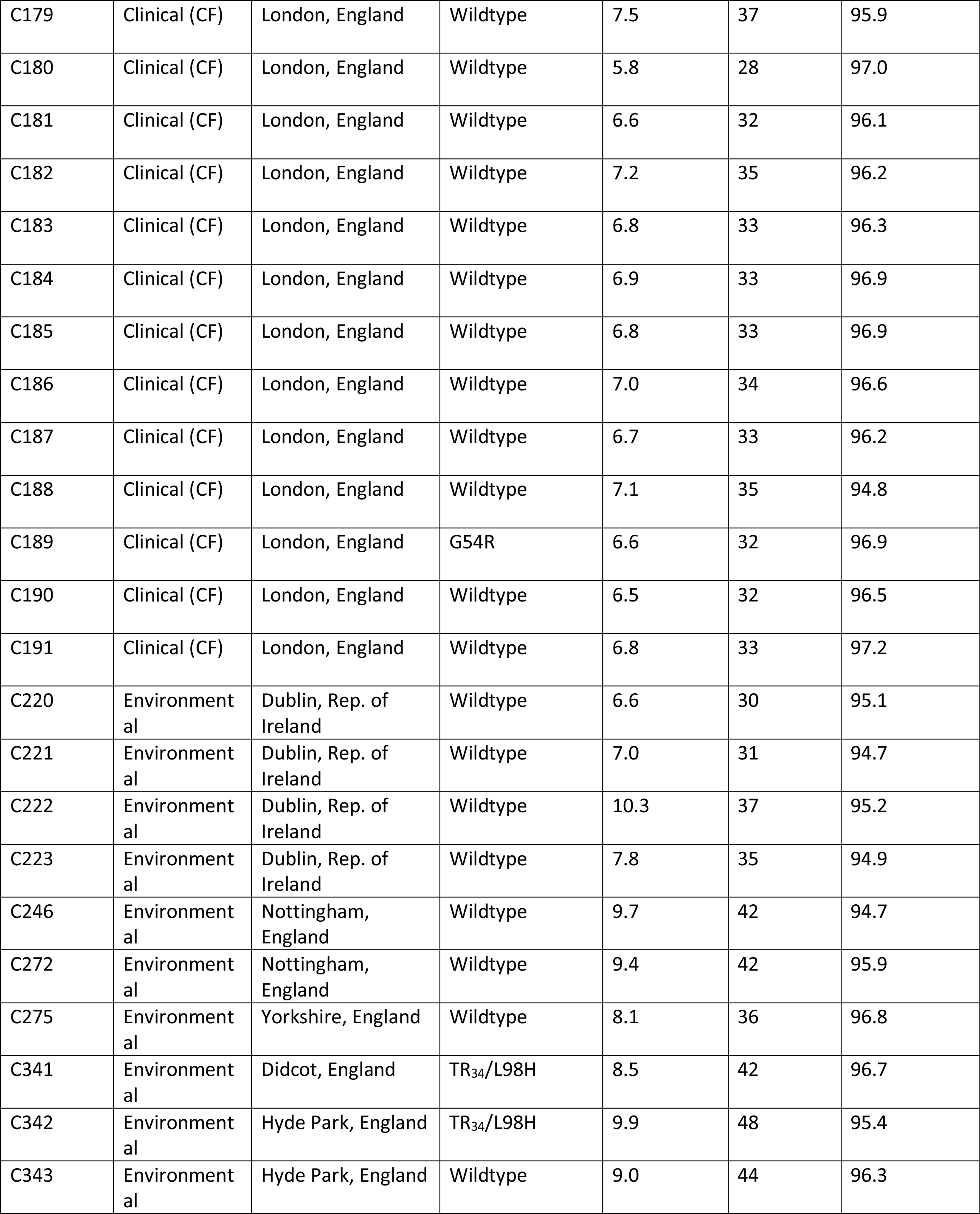
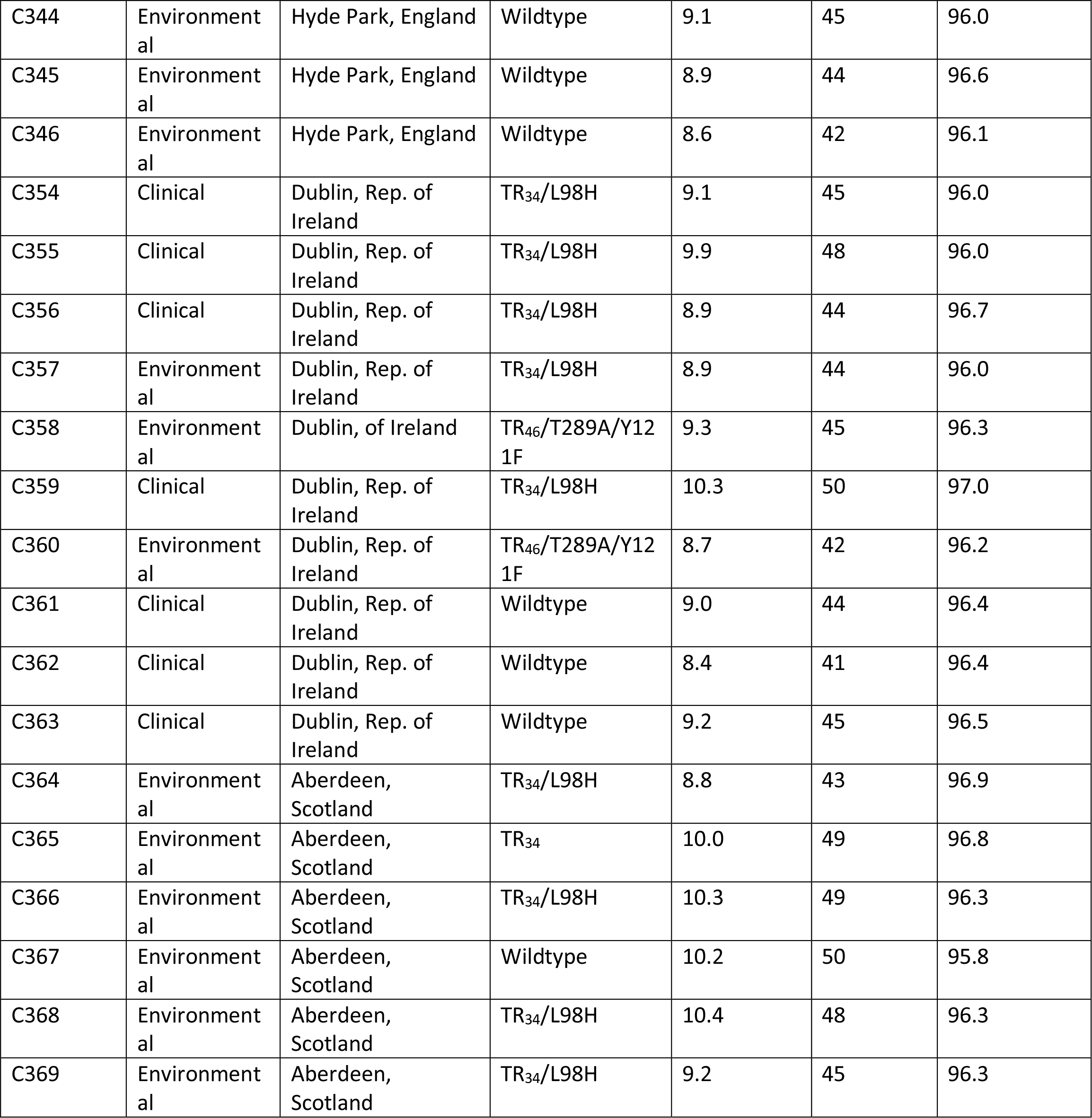
Clinical and environmental isolates of *A. fumigatus* used in this study, and details of whole-genome alignments. Patient cohort details for clinical isolates is provided, when available. Location information provided, where available. More accurate latitude and longitude details provided in the Microreact project. CF = cystic fibrosis; COPD = chronic obstructive pulmonary disease; RLL = right lower lobe; ABPA = allergic bronchopulmonary aspergillosis.

**Table S2.** In vitro antifungal susceptibility profiles of *A. fumigatus* isolates and corresponding resistance markers in *cyp51A*, mating type idiomorph and Clade membership as detected by whole genome sequencing. Minimum Inhibitory Concentrations above EUCAST (2018) clinical breakpoints, therefore indicating resistance, are shaded in grey. ND = not determined.

| Isolate | MIC (mg/litre) of<sup>a</sup>: |  |  | Resistance marker | Matin g type | Clade membershi p |
| --- | --- | --- | --- | --- | --- | --- |
|  | ITC | VOR | POS |  |  |  |
| ARAF001 | >16 | 1 | 0.5 | TR<sub>34</sub>/L98H | MAT1-1 | A |
| ARAF002 | >16 | 1 | 0.5 | TR<sub>34</sub>/L98H | MAT1-1 | A |
| ARAF003 | 1 | 0.25 | 0.06 | Wildtype | MAT1-2 | B |
| ARAF004 | 1 | 0.5 | 0.25 | Wildtype | MAT1-1 | A |
| ARAF005 | 0.5 | 0.125 | 0.06 | Wildtype | MAT1-2 | B |
| ARAF006 | 0.5 | 0.5 | 0.06 | Wildtype | MAT1-1 | A |
| C1 | >16 | 2 | 0.12 | TR<sub>34</sub>/L98H | MAT1-2 | A |
| C2 | >16 | 2 | 0.25 | TR<sub>34</sub>/L98H | MAT1-2 | A |
| C3 | >16 | 2 | 0.12 | TR<sub>34</sub>/L98H | MAT1-2 | A |
| C4 | >16 | 8 | <0.5 | Wildtype | MAT1-2 | B |
| C5 | >16 | 2 | 0.12 | TR<sub>34</sub>/L98H | MAT1-2 | A |
| C6 | >16 | 2 | 0.25 | TR<sub>34</sub>/L98H | MAT1-2 | A |
| C7 | >16 | 4 | 0.25 | TR<sub>34</sub>/L98H | MAT1-2 | A |
| C8 | >16 | 2 | 0.12 | TR<sub>34</sub>/L98H | MAT1-2 | A |
| C10 | >16 | 8 | 0.5 | TR<sub>34</sub>/L98H | MAT1-2 | A |
| C11 | >16 | 2 | 0.25 | TR<sub>34</sub>/L98H | MAT1-2 | A |
| C12 | >16 | 4 | 0.25 | TR<sub>34</sub>/L98H | MAT1-2 | A |
| C13 | >16 | 2 | 0.12 | TR<sub>34</sub>/L98H | MAT1-2 | A |
| C14 | >16 | >8 | 0.5 | TR<sub>34</sub>/L98H | MAT1-2 | A |
| C15 | >16 | 8 | <0.5 | TR<sub>34</sub>/L98H | MAT1-2 | A |
| C16 | >16 | 2 | 0.12 | TR<sub>34</sub>/L98H | MAT1-2 | A |
| C17 | >16 | 2 | 0.12 | TR<sub>34</sub>/L98H | MAT1-2 | A |
| C18 | >16 | 2 | 0.12 | TR<sub>34</sub>/L98H | MAT1-2 | A |
| C19 | >16 | 1 | 0.25 | TR<sub>34</sub>/L98H | MAT1-2 | A |
| C20 | >16 | 2 | <0.25 | TR<sub>34</sub>/L98H | MAT1-2 | A |
| C21 | >16 | 2 | 0.025 | TR<sub>34</sub>/L98H | MAT1-2 | A |
| C22 | >16 | 2 | 0.12 | TR<sub>34</sub>/L98H | MAT1-2 | A |
| C23 | >16 | 2 | 0.12 | TR<sub>34</sub>/L98H | MAT1-2 | A |
| C24 | >16 | 2 | 0.12 | TR<sub>34</sub>/L98H | MAT1-2 | A |
| C25 | >16 | 4 | 0.25 | TR<sub>34</sub>/L98H | MAT1-2 | A |
| C26 | >16 | 0.25 | 1 | G54R | MAT1-2 | A |
| C27 | >16 | 1 | 0.25 | TR<sub>34</sub>/L98H | MAT1-2 | A |
| C28 | >16 | 4 | 0.25 | TR<sub>34</sub>/L98H | MAT1-1 | A |
| C29 | >16 | 4 | 0.5 | TR<sub>34</sub>/L98H | MAT1-2 | A |
| C30 | >16 | 2 | 0.25 | TR<sub>34</sub>/L98H | MAT1-2 | A |
| C31 | >16 | 2 | 0.12 | TR<sub>34</sub>/L98H | MAT1-2 | A |
| C32 | >16 | 0.5 | 0.5 | TR<sub>34</sub>/L98H | MAT1-2 | A |
| C33 | >16 | 1 | 0.5 | TR<sub>34</sub>/L98H | MAT1-2 | A |
| C34 | <0.03 | 0.06 | 0.03 | Wildtype | MAT1-1 | B |
| C35 | 0.06 | 0.06 | 0.06 | Wildtype | MAT1-1 | A |
| C36 | 0.06 | 0.25 | 0.06 | Wildtype | MAT1-1 | A |
| C37 | 0.25 | 0.125 | 0.06 | Wildtype | MAT1-1 | B |
| C38 | 0.125 | 0.25 | 0.06 | Wildtype | MAT1-1 | A |
| C39 | 0.06 | 0.25 | 0.06 | Wildtype | MAT1-2 | B |
| C40 | 0.06 | 0.25 | 0.03 | Wildtype | MAT1-1 | B |
| C41 | 0.25 | 0.25 | 0.25 | Wildtype | MAT1-1 | B |
| C42 | 0.25 | 0.25 | 0.12 | Wildtype | MAT1-2 | B |
| C43 | 0.03 | 0.12 | 0.015 | Wildtype | MAT1-2 | B |
| C44 | 0.03 | 0.12 | 0.015 | Wildtype | MAT1-2 | B |
| C45 | 0.03 | 0.12 | 0.015 | Wildtype | MAT1-2 | B |
| C46 | 0.03 | 0.12 | 0.015 | Wildtype | MAT1-2 | B |
| C47 | 0.03 | 0.015 | 0.015 | Wildtype | MAT1-2 | B |
| C48 | 0.03 | 0.015 | 0.015 | Wildtype | MAT1-2 | B |
| C49 | 0.12 | 0.25 | 0.06 | Wildtype | MAT1-1 | B |
| C50 | 0.25 | 0.12 | 0.06 | Wildtype | MAT1-1 | B |
| C51 | 0.06 | 0.25 | 0.03 | Wildtype | MAT1-2 | B |
| C52 | <0.015 | 0.12 | 0.03 | Wildtype | MAT1-2 | B |
| C53 | 0.25 | 0.12 | 0.06 | Wildtype | MAT1-2 | B |
| C54 | 0.06 | 0.25 | 0.03 | Wildtype | MAT1-2 | B |
| C55 | 0.03 | 0.25 | 0.015 | Wildtype | MAT1-2 | B |
| C56 | 0.12 | 0.12 | 0.015 | Wildtype | MAT1-1 | B |
| C57 | 0.06 | 0.06 | 0.03 | Wildtype | MAT1-1 | B |
| C58 | 0.03 | 0.12 | 0.015 | Wildtype | MAT1-1 | A |
| C59 | 0.25 | 0.12 | 0.06 | Wildtype | MAT1-1 | A |
| C60 | 0.12 | 0.12 | 0.03 | Wildtype | MAT1-1 | A |
| C61 | 0.03 | 0.25 | 0.015 | Wildtype | MAT1-1 | A |
| C62 | 0.03 | 0.25 | 0.015 | Wildtype | MAT1-1 | A |
| C63 | 0.03 | 0.12 | 0.015 | Wildtype | MAT1-1 | A |
| C64 | 0.25 | 0.25 | 0.12 | Wildtype | MAT1-1 | A |
| C65 | 0.03 | 0.12 | 0.015 | Wildtype | MAT1-1 | A |
| C66 | 0.03 | 0.12 | 0.015 | Wildtype | MAT1-1 | A |
| C67 | 0.03 | 0.25 | 0.015 | Wildtype | MAT1-2 | B |
| C68 | 0.25 | 0.25 | 0.06 | TR<sub>34</sub>/L98H | MAT1-2 | A |
| C69 | 1 | 0.5 | 0.25 | TR<sub>34</sub>/L98H | MAT1-2 | A |
| C70 | 1 | 1 | 0.25 | TR<sub>34</sub>/L98H | MAT1-2 | A |
| C71 | 0.12 | 0.03 | 0.008 | Wildtype | MAT1-1 | B |
| C72 | 0.03 | 0.12 | 0.015 | Wildtype | MAT1-1 | B |
| C73 | 0.03 | 0.12 | 0.015 | Wildtype | MAT1-2 | B |
| C74 | 0.03 | 0.12 | 0.015 | Wildtype | MAT1-2 | B |
| C75 | 0.25 | 0.25 | 0.12 | Wildtype | MAT1-1 | A |
| C76 | 0.03 | 0.12 | 0.015 | Wildtype | MAT1-2 | B |
| C77 | 0.06 | 0.25 | 0.03 | Wildtype | MAT1-2 | B |
| C78 | ND | ND | ND | TR<sub>34</sub>/L98H | MAT1-1 | A |
| C79 | ND | ND | ND | TR<sub>46</sub>/Y121F/T289A | MAT1-2 | A |
| C80 | ND | ND | ND | TR<sub>34</sub>/L98H | MAT1-2 | A |
| C81 | 0.25 | 0.25 | 0.06 | Wildtype | MAT1-1 | B |
| C82 | 4 | 2 | 1 | TR<sub>34</sub>/L98H | MAT1-2 | A |
| C83 | 16 | 2 | 0.5 | TR<sub>34</sub>/L98H | MAT1-2 | A |
| C84 | 4 | 4 | 0.5 | TR<sub>34</sub>/L98H | MAT1-2 | A |
| C85 | ND | ND | ND | TR<sub>34</sub>/L98H | MAT1-2 | A |
| C86 | ND | ND | ND | TR<sub>34</sub>/L98H | MAT1-1 | A |
| C87 | ND | ND | ND | TR<sub>46</sub>/Y121F/T289A | MAT1-1 | B |
| C88 | ND | ND | ND | TR<sub>34</sub>/L98H | MAT1-2 | A |
| C89 | ND | ND | ND | TR<sub>46</sub>/Y121F/T289A | MAT1-2 | A |
| C91 | ND | ND | ND | TR<sub>46</sub>/Y121F/T289A | MAT1-1 | B |
| C92 | ND | ND | ND | TR<sub>34</sub>/L98H | MAT1-2 | A |
| C93 | ND | ND | ND | TR<sub>46</sub>/Y121F/T289A | MAT1-2 | A |
| C95 | ND | ND | ND | Wildtype | MAT1-1 | B |
| C96 | ND | ND | ND | Wildtype | MAT1-2 | B |
| C99 | 2 | 2 | 0.5 | TR<sub>34</sub>/L98H | MAT1-1 | A |
| C100 | 16 | 4 | ND | TR<sub>34</sub>/L98H | MAT1-2 | A |
| C103 | 1 | 2 | ND | TR<sub>34</sub>/L98H | MAT1-2 | A |
| C104 | >16 | 0.5 | 1 | TR<sub>34</sub>/L98H | MAT1-1 | A |
| C105 | >16 | 4 | ND | TR<sub>34</sub>/L98H | MAT1-2 | A |
| C106 | 2 | 4 | ND | TR<sub>34</sub>/L98H | MAT1-2 | A |
| C107 | 1 | 4 | ND | TR<sub>34</sub>/L98H | MAT1-2 | A |
| C108 | 16 | 1 | ND | TR<sub>34</sub>/L98H | MAT1-2 | A |
| C109 | >16 | 1 | ND | TR<sub>34</sub>/L98H | MAT1-1 | B |
| C110 | >16 | 1 | ND | TR<sub>34</sub>/L98H | MAT1-2 | A |
| C111 | 16 | 2 | ND | TR<sub>34</sub>/L98H | MAT1-1 | A |
| C112 | 16 | 2 | ND | TR<sub>34</sub>/L98H | MAT1-2 | A |
| C113 | ND | 0.25 | ND | Wildtype | MAT1-1 | B |
| C114 | >16 | 2 | ND | TR<sub>34</sub>/L98H | MAT1-1 | A |
| C115 | 4 | 1 | 0.25 | TR<sub>34</sub>/L98H | MAT1-1 | A |
| C117 | 1 | 2 | ND | Wildtype | MAT1-2 | B |
| C118 | 8 | 1 | ND | Wildtype | MAT1-2 | B |
| C119 | 16 | 1 | ND | TR<sub>34</sub>/L98H | MAT1-2 | A |
| C120 | >16 | 0.125 | ND | G54R | MAT1-1 | B |
| C121 | >16 | 0.25 | 0.5 | G54E | MAT1-1 | A |
| C122 | >16 | 2 | ND | TR<sub>34</sub>/L98H | MAT1-2 | A |
| C123 | >16 | 2 | 0.25 | TR<sub>34</sub>/L98H | MAT1-2 | A |
| C124 | >16 | 0.125 | 4 | G54W | MAT1-1 | B |
| C125 | >16 | 2 | 2 | TR<sub>34</sub>/L98H | MAT1-2 | A |
| C126 | >16 | 0.03 | 2 | G54W | MAT1-1 | B |
| C127 | >16 | 0.25 | 0.125 | Wildtype | MAT1-2 | B |
| C128 | >16 | 2 | 1 | Wildtype | MAT1-2 | B |
| C129 | 8 | 2 | 0.5 | Wildtype | MAT1-2 | B |
| C130 | >16 | 0.125 | 8 | G54W | MAT1-2 | B |
| C131 | >16 | 0.125 | 16 | G54W | MAT1-2 | B |
| C132 | >16 | 0.125 | 8 | G54W | MAT1-2 | B |
| C133 | >16 | 2 | 2 | TR<sub>34</sub>/L98H | MAT1-2 | A |
| C134 | 16 | 2 | 1 | TR<sub>34</sub>/L98H | MAT1-1 | B |
| C135 | >16 | 0.125 | 16 | G54W | MAT1-2 | B |
| C136 | >16 | 2 | 2 | Wildtype | MAT1-2 | B |
| C137 | >16 | 2 | 2 | Wildtype | MAT1-2 | B |
| C138 | >16 | 0.25 | 2 | P216L | MAT1-1 | B |
| C139 | >16 | 2 | 1 | Wildtype | MAT1-2 | B |
| C140 | >16 | 0.125 | 16 | G54W | MAT1-1 | B |
| C141 | 16 | 1 | 0.25 | TR<sub>34</sub>/L98H | MAT1-1 | B |
| C142 | >16 | 0.06 | 0.25 | G54W | MAT1-1 | A |
| C143 | 16 | 2 | 0.5 | TR<sub>34</sub>/L98H | MAT1-2 | A |
| C144 | >16 | 2 | 0.25 | TR<sub>34</sub>/L98H | MAT1-2 | A |
| C145 | >16 | 0.125 | 0.5 | P216L | MAT1-2 | B |
| C146 | 0.5 | 0.125 | 0.03 | Wildtype | MAT1-2 | B |
| C147 | 1 | 0.125 | 0.03 | Wildtype | MAT1-2 | B |
| C148 | 2 | 0.125 | 0.03 | Wildtype | MAT1-2 | B |
| C149 | >16 | 0.25 | 0.125 | TR<sub>34</sub>/L98H | MAT1-1 | A |
| C150 | 16 | 0.25 | 0.125 | TR<sub>34</sub>/L98H | MAT1-1 | A |
| C151 | >16 | 0.25 | 0.125 | TR<sub>34</sub>/L98H | MAT1-1 | A |
| C152 | 0.5 | 0.125 | 0.03 | Wildtype | MAT1-2 | B |
| C153 | 1 | 0.125 | 0.03 | Wildtype | MAT1-2 | B |
| C154 | 1 | 0.125 | 0.03 | Wildtype | MAT1-2 | B |
| C155 | 16 | >16 | 4 | TR<sub>34</sub>/L98H/T289A/I364V/G448S | MAT1-2 | A |
| C156 | 16 | >16 | 4 | TR<sub>34</sub>/L98H/T289A/I364V/G448S | MAT1-2 | A |
| C157 | 16 | >16 | 4 | TR<sub>34</sub>/L98H/T289A/I364V/G448S | MAT1-2 | A |
| C158 | 16 | >16 | 4 | TR<sub>34</sub>/L98H/T289A/I364V/G448S | MAT1-2 | A |
| C159 | >16 | 2 | 0.5 | TR<sub>34</sub>/L98H | MAT1-2 | A |
| C160 | >16 | 2 | 0.5 | TR<sub>34</sub>/L98H | MAT1-2 | A |
| C161 | 4 | 0.5 | 0.125 | TR<sub>34</sub>/L98H | MAT1-1 | A |
| C162 | 2 | 2 | 1 | Wildtype | MAT1-1 | A |
| C163 | >16 | 1 | >16 | G54W | MAT1-1 | A |
| C164 | 2 | 1 | 0.125 | Wildtype | MAT1-2 | B |
| C165 | 2 | 0.5 | 0.125 | Wildtype | MAT1-1 | B |
| C166 | 1 | 0.25 | 0.06 | Wildtype | MAT1-2 | B |
| C167 | 0.06 | 0.06 | 0.03 | Wildtype | MAT1-1 | B |
| C168 | 2 | 0.25 | 0.03 | Wildtype | MAT1-1 | B |
| C169 | 0.06 | 0.06 | 0.015 | Wildtype | MAT1-2 | B |
| C170 | 0.06 | 0.06 | 0.015 | Wildtype | MAT1-2 | B |
| C171 | 16 | 0.5 | 0.25 | TR<sub>34</sub>/L98H | MAT1-2 | A |
| C172 | 0.06 | 0.06 | 0.03 | Wildtype | MAT1-2 | B |
| C173 | 0.06 | 0.06 | 0.015 | Wildtype | MAT1-1 | B |
| C174 | 0.06 | 0.06 | 0.03 | Wildtype | MAT1-1 | B |
| C175 | 0.06 | 0.06 | 0.03 | Wildtype | MAT1-2 | B |
| C176 | 0.06 | 0.06 | 0.03 | Wildtype | MAT1-1 | B |
| C177 | 0.06 | 0.125 | 0.03 | Wildtype | MAT1-1 | A |
| C178 | 0.06 | 0.06 | 0.03 | Wildtype | MAT1-2 | B |
| C179 | 0.06 | 0.06 | 0.03 | Wildtype | MAT1-1 | B |
| C180 | 0.06 | 0.06 | 0.03 | Wildtype | MAT1-2 | B |
| C181 | 0.06 | 0.125 | 0.06 | Wildtype | MAT1-1 | B |
| C182 | 0.125 | 0.125 | 0.03 | Wildtype | MAT1-2 | B |
| C183 | 0.125 | 0.06 | 0.03 | Wildtype | MAT1-2 | B |
| C184 | 0.06 | 0.06 | 0.03 | Wildtype | MAT1-2 | B |
| C185 | 0.06 | 0.06 | 0.03 | Wildtype | MAT1-1 | B |
| C186 | 0.06 | 0.06 | 0.03 | Wildtype | MAT1-1 | A |
| C187 | 0.06 | 0.06 | 0.03 | Wildtype | MAT1-2 | B |
| C188 | 0.125 | 0.06 | 0.03 | Wildtype | MAT1-1 | A |
| C189 | >16 | 0.06 | 0.25 | G54R | MAT1-2 | B |
| C190 | 0.06 | 0.03 | 0.03 | Wildtype | MAT1-1 | B |
| C191 | 0.06 | 0.06 | 0.015 | Wildtype | MAT1-2 | B |
| C220 | 0.06 | 0.06 | 0.06 | Wildtype | MAT1-1 | A |
| C221 | 0.125 | 0.06 | 0.06 | Wildtype | MAT1-2 | A |
| C222 | 0.06 | 0.06 | 0.03 | Wildtype | MAT1-2 | B |
| C223 | 0.125 | 0.06 | 0.06 | Wildtype | MAT1-1 | A |
| C246 | 0.125 | 0.125 | 0.06 | Wildtype | MAT1-2 | A |
| C272 | 0.125 | 0.125 | 0.03 | Wildtype | MAT1-2 | B |
| C275 | 0.125 | 0.125 | 0.06 | Wildtype | MAT1-2 | B |
| C341 | >16 | 0.5 | 0.25 | TR<sub>34</sub>/L98H | MAT1-1 | A |
| C342 | >16 | 0.5 | 0.25 | TR<sub>34</sub>/L98H | MAT1-2 | A |
| C343 | 0.06 | 0.06 | 0.03 | Wildtype | MAT1-2 | B |
| C344 | 0.125 | 0.06 | 0.03 | Wildtype | MAT1-1 | B |
| C345 | 0.25 | 0.06 | 0.03 | Wildtype | MAT1-2 | B |
| C356 | 0.25 | 0.06 | 0.03 | Wildtype | MAT1-2 | B |
| C354 | 4 | 2 | 0.5 | TR<sub>34</sub>/L98H | MAT1-2 | A |
| C355 | 2 | 2 | 0.5 | TR<sub>34</sub>/L98H | MAT1-1 | A |
| C356 | 0.5 | 2 | 0.5 | TR<sub>34</sub>/L98H | MAT1-2 | A |
| C357 | ND | ND | ND | TR<sub>34</sub>/L98H | MAT1-1 | A |
| C358 | ND | ND | ND | TR<sub>46</sub>/Y121F/T289A | MAT1-2 | A |
| C359 | 32 | 2 | 0.5 | TR<sub>34</sub>/L98H | MAT1-2 | A |
| C360 | ND | ND | ND | TR<sub>46</sub>/Y121F/T289A | MAT1-2 | A |
| C361 | ND | ND | ND | Wildtype | MAT1-2 | A |
| C362 | 0.75 | 0.125 | 0.25 | Wildtype | MAT1-2 | A |
| C363 | 0.03 | 0.12 | 0.25 | Wildtype | MAT1-2 | A |
| C364 | 4 | 0.5 | ND | TR<sub>34</sub>/L98H | MAT1-1 | A |
| C365 | 16 | 2 | ND | TR<sub>34</sub> | MAT1-1 | A |
| C366 | 4 | 0.25 | ND | TR<sub>34</sub>/L98H | MAT1-1 | A |
| C367 | 1 | 0.5 | ND | Wildtype | MAT1-2 | A |
| C368 | 4 | 0.5 | ND | TR<sub>34</sub>/L98H | MAT1-2 | A |
| C369 | 1 | 0.5 | ND | TR<sub>34</sub>/L98H | MAT1-1 | A |

**Table S3.** Average value of Tajima’s *D* statistic per chromosome for isolates within Clades A and B, and average and F*_ST_* values per chromosome. Range of values are shown in brackets.

| Chromosome | Clade A | Clade B | *F<sub>ST</sub>* |
| --- | --- | --- | --- |
| I | 0.4047 (-1.9766-3.9560) | -0.5535 (-2.6785-2.6761) | 0.1022 (0-0.9440) |
| II | 0.5241 (-2.4022-5.5133) | -0.3028 (-2.0764-4.8010) | 0.1156 (0.0069-0.3539) |
| III | 0.3267 (-2.1441-4.3288) | -0.3422 (-2.5142-3.2920) | 0.1273 (0.0092-0.4244) |
| IV | 0.2774 (-2.1356-3.9701) | -0.4883 (-2.4272-3.7844) | 0.1622 (0.0115-0.5633) |
| V | 0.3775 (-2.26076-4.3397) | -0.3317 (-2.2117-3.4575) | 0.1616 (0.0168-0.5171) |
| VI | 0.7591 (-2.0187-4.1077) | -0.5145 (-2.5173-3.6985) | 0.1055 (0.0131-0.4314) |
| VII | 0.7888 (-2.0023-4.1684) | -0.2754 (-2.6883-3.8941) | 0.1530 (0.0208-0.5591) |
| VIII | 0.6067 (-2.0557-4.4425) | -0.0396 (-2.2204-3.1474) | 0.1530 (0.0202-0.3710) |
| Mitochondria (MT) | 0.5311 (-1.0274-2.6496) | -0.7255 (-0.9931- -0.2730) | 0.1520 (0.0682-0.2765) |

**Table S4.** Genes found within regions of high F*_ST_* in Chromosome 1 when performing fixation index analysis between Clades A and B in non-overlapping windows of 10 kb

| Gene ID | Genomic Location | Product Description |
| --- | --- | --- |
| Afu1g15000 | 4,029,090 – 4,031,693 | Putative isopropylmalate synthase |
| Afu1g15010 | 4,031,939 – 4,038,777 | Ortholog of *A. flavus* putative AMP binding domain protein AFLA_084630 |
| Afu1g15020 | 4,039,173 – 4,040,722 | 40S ribosomal protein S5 |
| Afu1g16860 | 4,598,911 – 4,600,377 | Ortholog of *A. flavus* conserved hypothetical protein ALFA_082490 |
| Afu1g16870 | 4,600,954 – 4,602,774 | Ortholog of *A. flavus* conserved hypothetical protein AFLA_016260 |
| Afu1g16880 | 4,603,153 – 4,608,184 | Ortholog of *A. flavus* putative ABC multidrug transporter AFLA_082400 |
| Afu1g16890 | 4,608,535 – 4,609,830 | Ortholog of *A. flavus* putative transesterase LovD AFLA_063440 |
| Afu1g16900 | 4,615,099 – 4,617,349 | Ortholog of *A. fumigatus* A1163<br>AFUB_016290 (protein of unknown function) |
| Afu1g16910 | 4,617,552 – 4,619,374 | Has domain with predicted role in<br>transmembrane transport |

**Table S5.** Pairs or groups of *A. fumigatus* isolates from both environmental and clinical sources with high genetic relatedness.

| Pair/<br>Group | Isolates<br>included<br>(Clade) | Mating<br>idiomorph | Source | *cyp51A* allele | Bootstrap<br>support | Average<br>SNPs<br>separating<br>isolates | t-test *p*-<br>value |
| --- | --- | --- | --- | --- | --- | --- | --- |
| 1 | C21 (A<sub>A</sub>)<br>C112 (A<sub>A</sub>) | MAT1-2<br>MAT1-2 | Environmental<br>Clinical | TR<sub>34</sub>/L98H<br>TR<sub>34</sub>/L98H | 100% | 227 | <3.0473<sup>e-252</sup> |
| 2 | C133 (A<sub>A</sub>)<br>C23 (A<sub>A</sub>) | MAT1-2<br>MAT1-2 | Clinical<br>Environmental | TR<sub>34</sub>/L98H<br>TR<sub>34</sub>/L98H | 65% | 270 | <3.0473<sup>e-252</sup> |
| 3 | C354 (A)<br>C355 (A)<br>C357 (A) | MAT1-2<br>MAT1-1<br>MAT1-1 | Clinical<br>Clinical<br>Environmental | TR<sub>34</sub>/L98H<br>TR<sub>34</sub>/L98H<br>TR<sub>34</sub>/L98H | 100% | 237 | <3.0473<sup>e-252</sup> |
| 4 | C191 (B)<br>C275 (B) | MAT1-2<br>MAT1-2 | Clinical<br>Environmental | Wildtype<br>Wildtype | 100% | 581 | <3.0473<sup>e-252</sup> |
| 5 | C96 (B)<br>C42 (B)<br>C43 (B)<br>C44 (B)<br>C48 (B) | MAT1-2<br>MAT1-2<br>MAT1-2<br>MAT1-2<br>MAT1-2 | Environmental<br>Clinical<br>Clinical<br>Clinical<br>Clinical | Wildtype<br>Wildtype<br>Wildtype<br>Wildtype<br>Wildtype | 79% | 217 | 3.0473<sup>e-252</sup> |
| 6 | C7 (A)<br>C25 (A)<br>C82 (A) | MAT1-2<br>MAT1-2<br>MAT1-2 | Environmental<br>Environmental<br>Clinical | TR<sub>34</sub>/L98H<br>TR<sub>34</sub>/L98H<br>TR<sub>34</sub>/L98H | 100% | 252 | 1.7857<sup>e-112</sup> |

### Supplementary Figures

**Figure S1:**
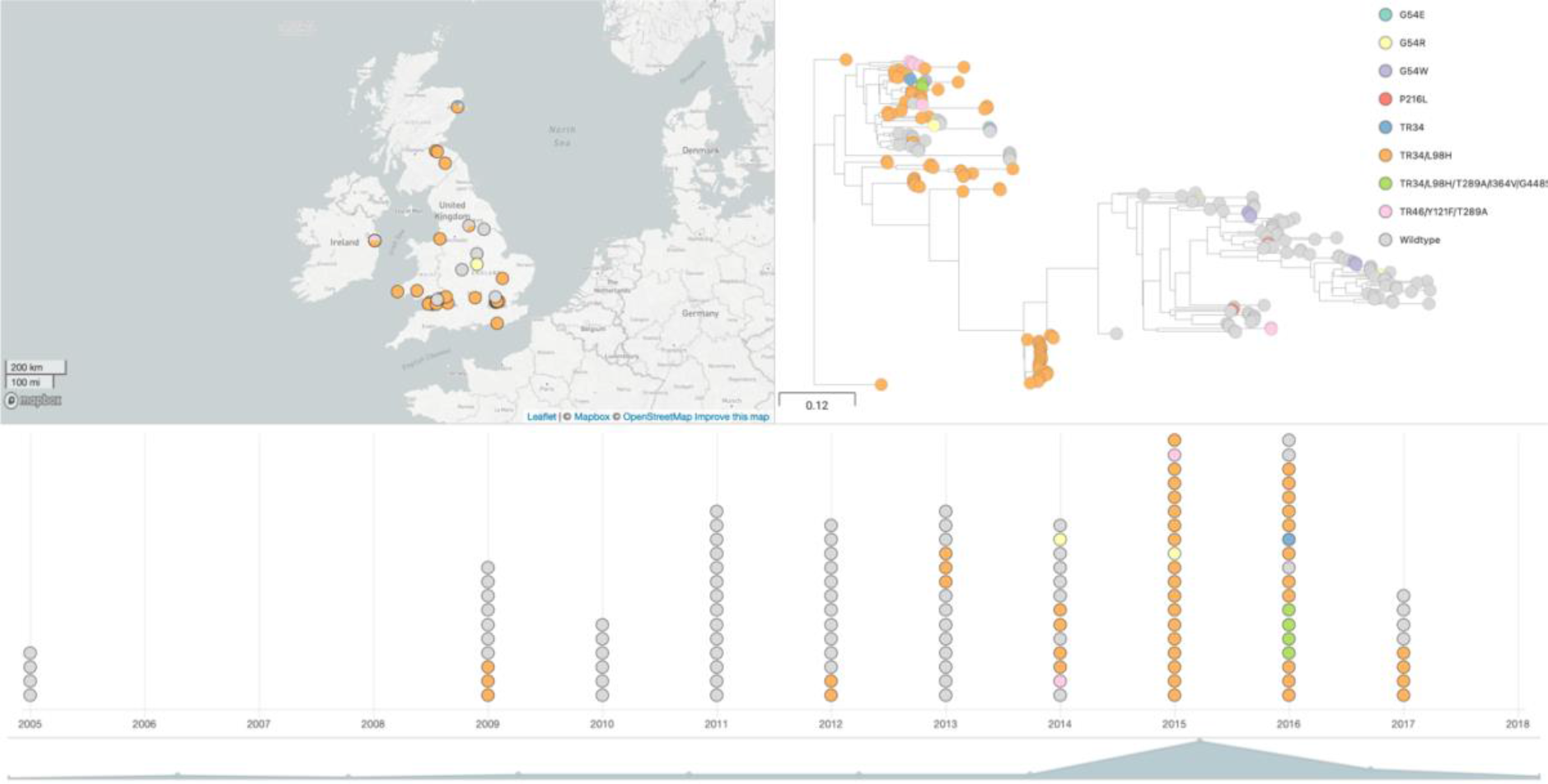
Microreact project screenshot of the dataset https://microreact.org/project/viUDBzrCmTNKmY9Fu6Zhxi

**Figure S2:**
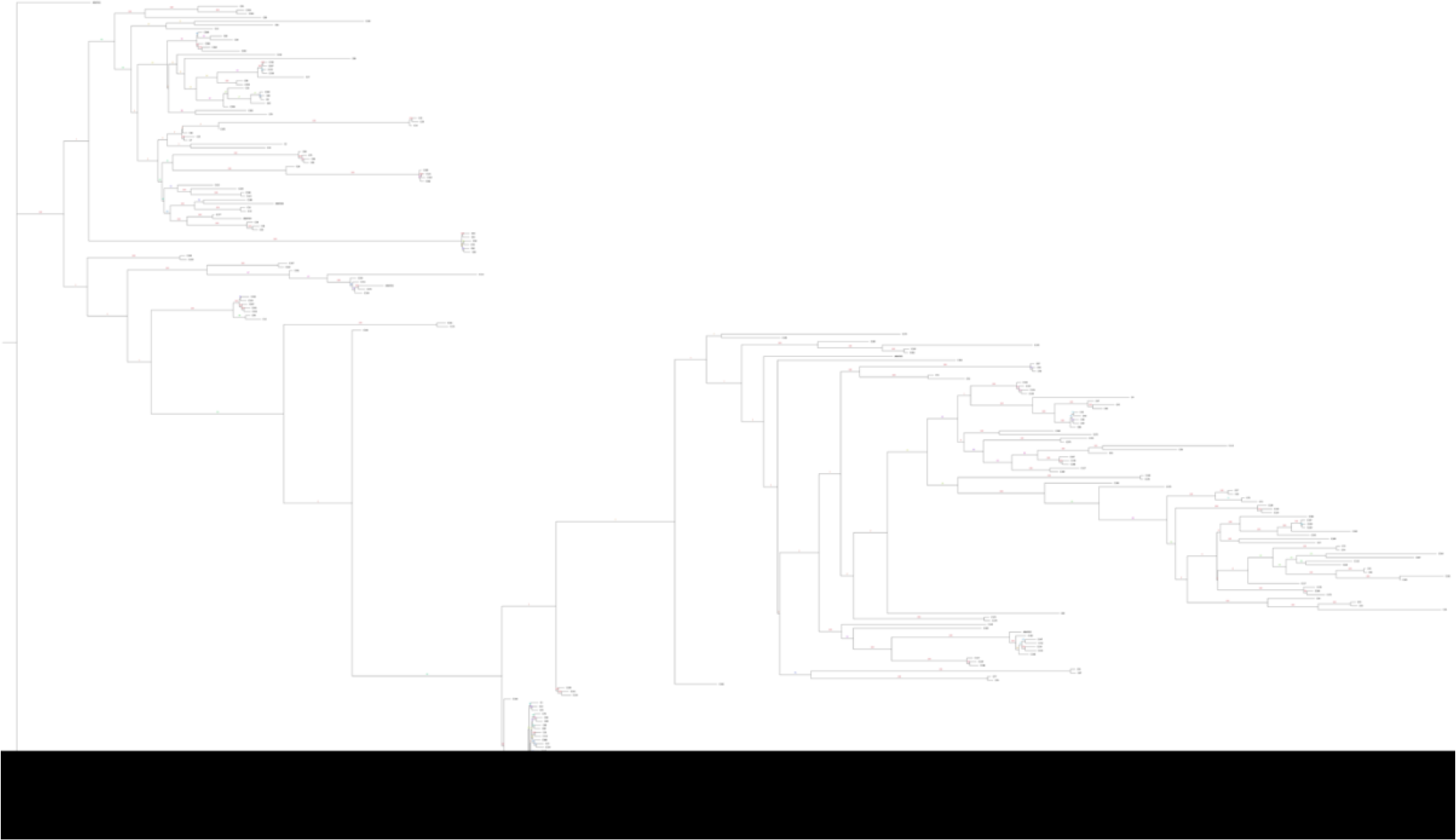
Phylogenetic analysis of all 218 *A. fumigatus* isolates with bootstrap support over 1000 replicates performed on WGS SNP data to generate maximum-likelihood phylogeny. Branch lengths represent average number of SNPs.

**Figure S3:**
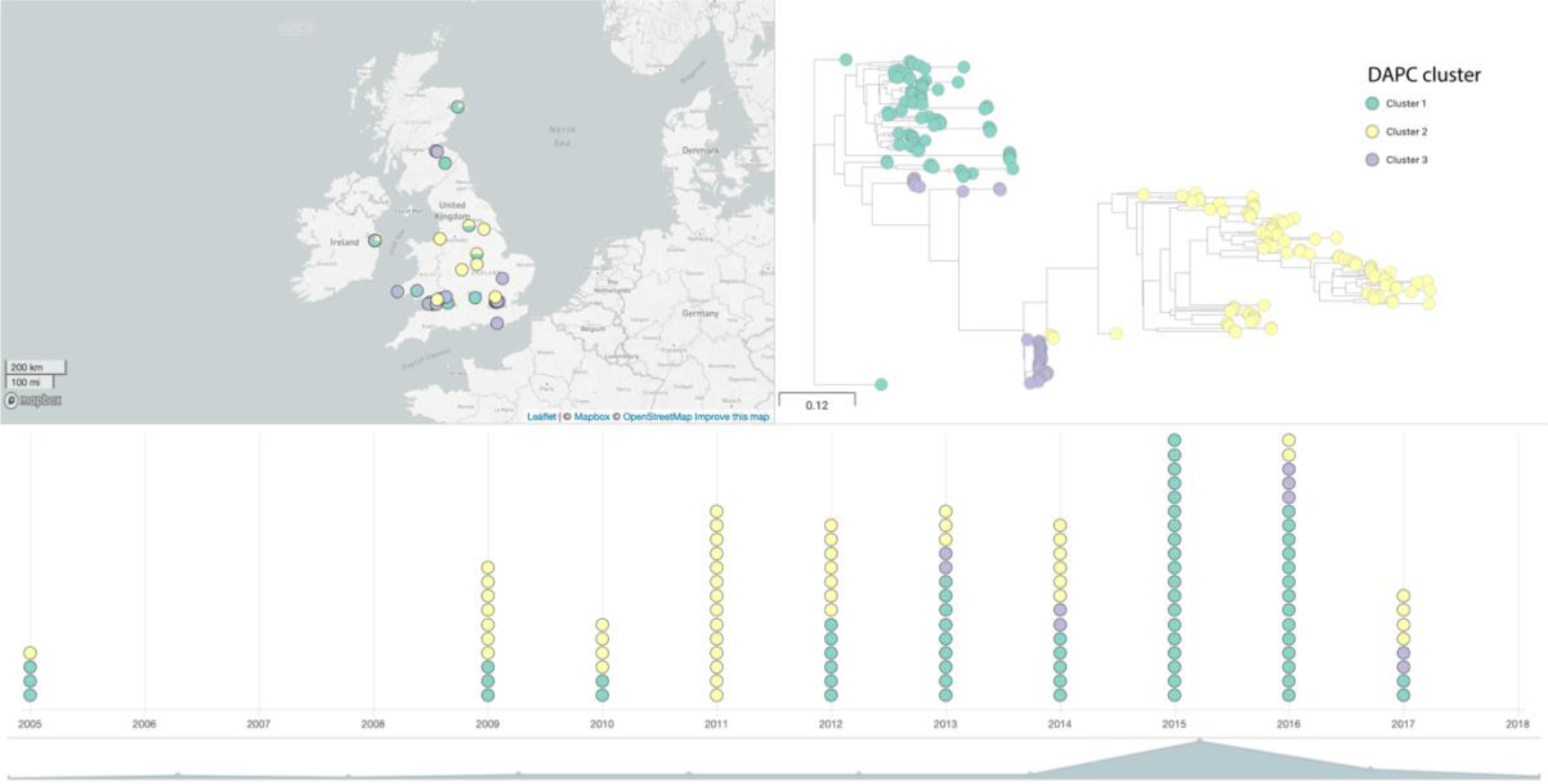
Microreact project screenshot of the dataset with DAPC clusters showing lack of geographic and temporal clustering.

**Figure S4:**
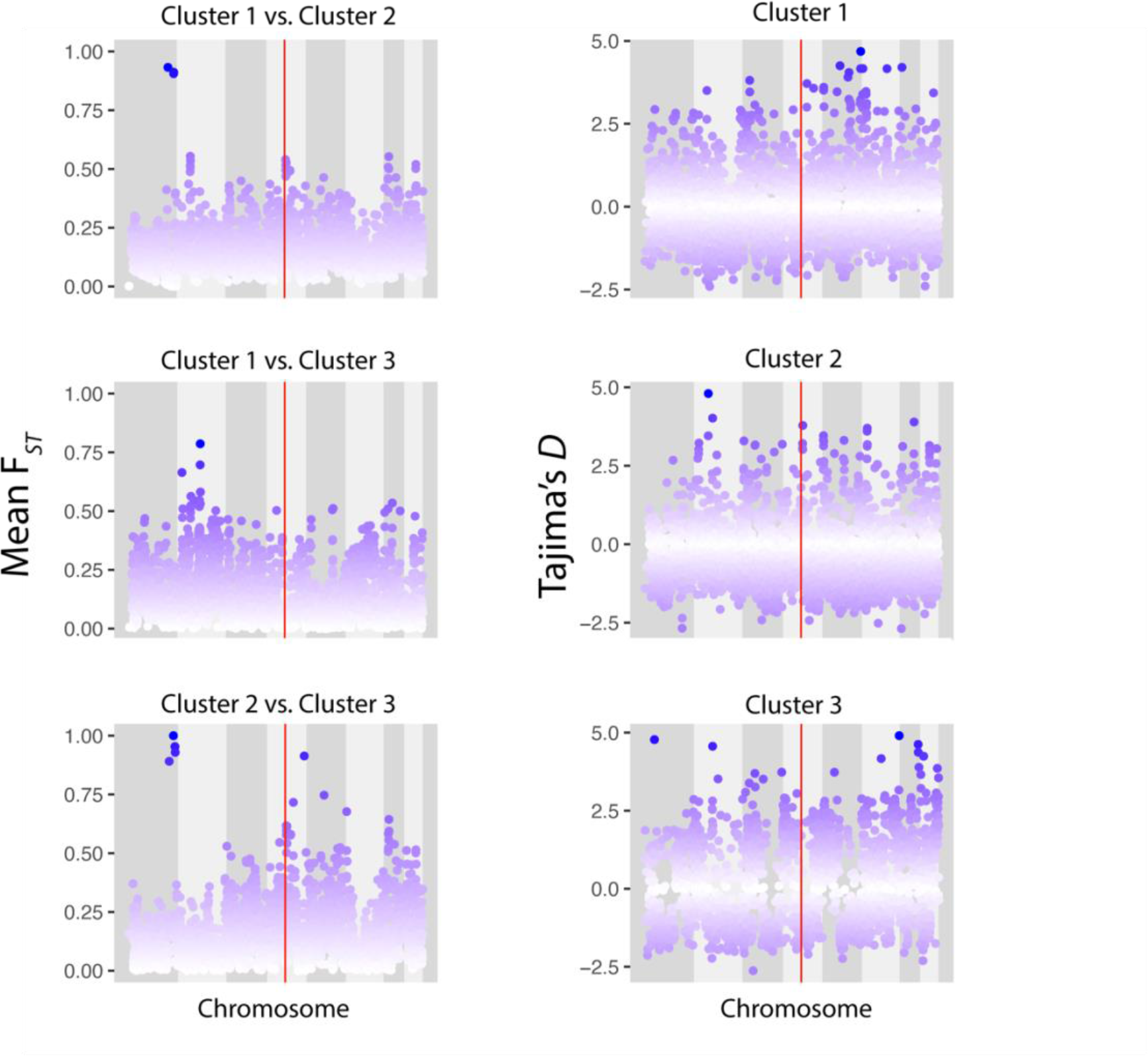
Scatterplots of sliding 10-kb non-overlapping window estimates of F*_ST_* for each chromosome between isolates within Clusters 1 and 2, Clusters 1 and 3, and Clusters 2 and 3 (top to bottom, left panel). Scatterplots of Tajima’s D estimates for each chromosome for all isolates within Clusters 1, 2 and 3 (right panel). The position of *cyp51A* is highlighted in red.

**Figure S5:**
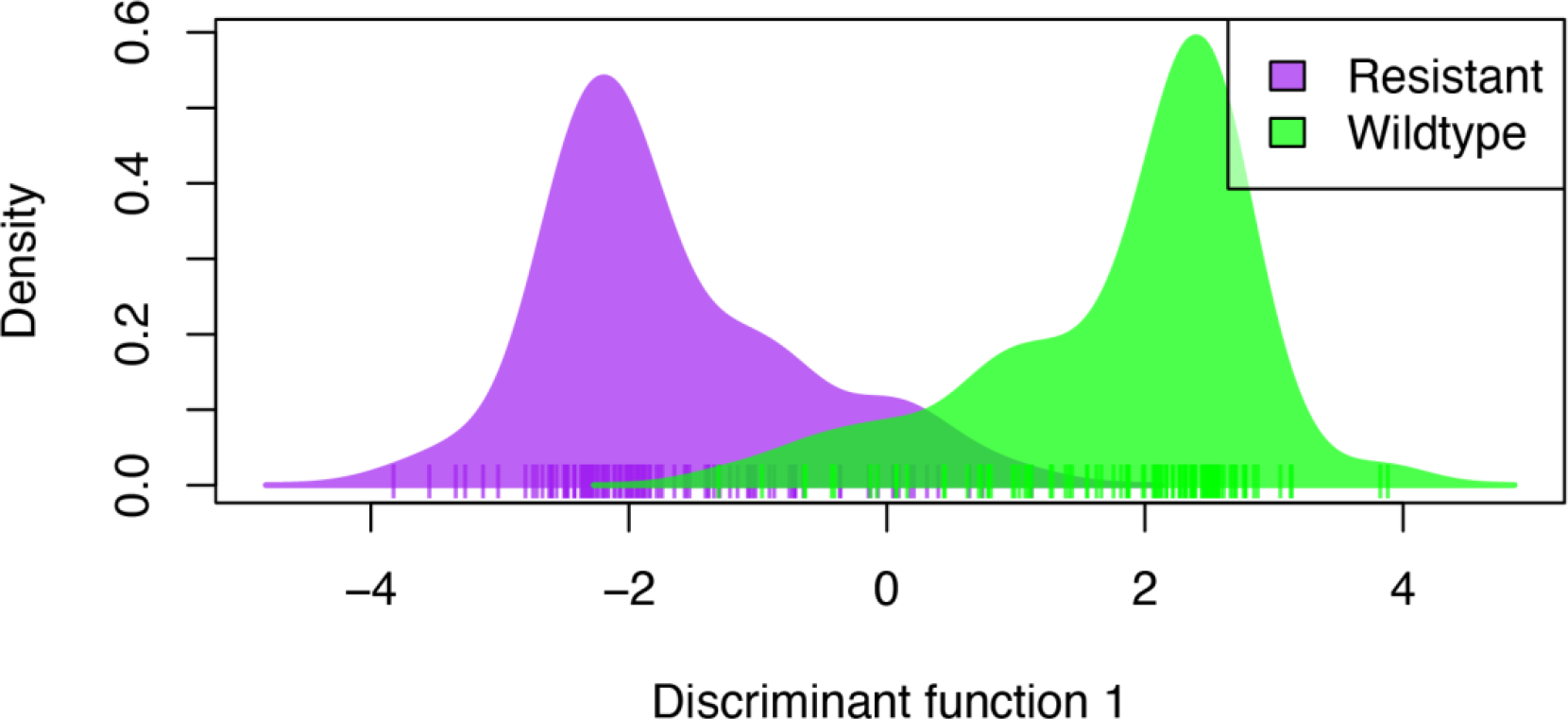
Azole-resistant and azole-sensitive *A. fumigatus* isolates form different, distinct clusters Discriminant Analysis of Principal Components (DAPC) on all azole-resistant and - sensitive (wildtype for *cyp51A*) *A. fumigatus* isolates using the first two principal components. This analysis illustrates slight genetic identity between azole-resistant and - sensitive isolates, with the overlap illustrating some similar genetic backgrounds have been observed.

### Supplementary Data

**Supplementary Data 1 (Excel format):** Significant SNPs for itraconazole resistance using the TreeWAS subsequent association test, in order of ascending p-value for all *A. fumigatus* isolates in this study with known itraconazole MICs. Chromosome, SNP position and gene loci listed along with gene description and type of SNP (synonymous or non-synonymous).

**Supplementary Data 2 (Excel format):** Significant SNPs for itraconazole resistance using the TreeWAS terminal association test, in order of ascending p-value for all *A. fumigatus* isolates in this study with known itraconazole MICs. Chromosome, SNP position and gene loci listed along with gene description and type of SNP (synonymous or non-synonymous).

**Supplementary Data 3 (Excel format):** Significant SNPs for itraconazole resistance using the TreeWAS simultaneous association test, in order of ascending p-value for all *A. fumigatus* isolates in this study with known itraconazole MICs. Chromosome, SNP position and gene loci listed along with gene description and type of SNP (synonymous or non-synonymous).

**Supplementary Data 4 (Excel format):** Summary of 184 genes found in region of high F*_ST_* in chromosome 1 when comparing clades A and B. Associated *p*-values from TreeWAS for genes containing SNPs that are statistically significant for itraconazole resistance are also included.

